# Long-chain polyphosphates impair SARS-CoV-2 infection and replication: a route for therapy in man

**DOI:** 10.1101/2020.11.18.388413

**Authors:** Veronica Ferrucci, Dae-Young Kong, Fatemeh Asadzadeh, Laura Marrone, Roberto Siciliano, Pellegrino Cerino, Giuseppina Criscuolo, Ida Pisano, Fabrizio Quarantelli, Barbara Izzo, Giovanna Fusco, Marika Comegna, Angelo Boccia, Maurizio Viscardi, Giorgia Borriello, Sergio Brandi, Bianca Maria Pierri, Claudia Tiberio, Luigi Atripaldi, Giovanni Paolella, Giuseppe Castaldo, Stefano Pascarella, Martina Bianchi, Rosa Della Monica, Lorenzo Chiariotti, Kyong-Seop Yun, Jae-Ho Cheong, Hong-Yeoul Kim, Massimo Zollo

## Abstract

Anti-viral activities of long-chain inorganic polyphosphates (PolyPs) against severe acute respiratory syndrome coronavirus (SARS-CoV)-2 infection were investigated. In molecular docking analyses, PolyPs interacted with several conserved angiotensin-converting enzyme (ACE)2 and RNA-dependent RNA polymerase (RdRp) amino acids. We thus tested PolyPs for functional interactions *in vitro* in SARS-CoV-2–infected Vero E6, Caco2 and human primary nasal epithelial cells. Immunofluorescence, qPCR, direct RNA sequencing, FISH and Immunoblotting were used to determine virus loads and transcription levels of genomic(g)RNAs and sub-genomic(sg)RNAs. We show that PolyP120 binds to ACE2 and enhances its proteasomal degradation. PolyP120 shows steric hindrance of the genomic Sars-CoV-2-RNA/RdRP complex, to impair synthesis of positive-sense gRNAs, viral subgenomic transcripts and structural proteins needed for viral replication. Thus, PolyP120 impairs infection and replication of Korean and European (containing non-synonymous variants) SARS-CoV-2 strains. As PolyPs have no toxic activities, we envision their use as a nebulised formula for oropharyngeal delivery to prevent infections of SARS-CoV-2 and during early phases of antiviral therapy.

## Introduction

Coronaviruses (CoVs) contain positive-sense, single-stranded RNA (~30 kb). Four major categories have been reported, with alphaCoV and betaCoV known to infect humans. Those that can replicate in the lower respiratory tract cause pneumonia, which can be fatal (1),(2), and these include severe acute respiratory syndrome (SARS)-CoV, Middle East respiratory syndrome (MERS)-CoV, and the new SARS-CoV-2. This last CoV belongs to the betaCoV genus (3) and has resulted in pandemic acute respiratory syndrome in humans (i.e., COVID-19 disease). This can progress to acute respiratory distress syndrome (ARDS), generally around 8 to 9 days after symptom onset (4). Like the other respiratory CoVs, SARS-CoV-2 is transmitted via respiratory droplets, with possible faecal–oral transmission (5).

When SARS-CoV-2 infects host cells, it replicates its genomic (g)RNA to produce smaller RNAs known as subgenomic (sg)RNAs, mostly used to synthesise conserved structural viral proteins Spike (S), Envelope (E), Membrane (M) and Nucleocapsid (N) (6).

The receptor used by SARS-CoV-2 to enter host cells is angiotensin-converting enzyme 2 (ACE2), which is mainly expressed in the lungs (7),(8), together with the cellular serine protease TMPRSS2 (9).

The SARS-CoV-2 RNA-dependent RNA polymerase (RdRp; also, non-structural protein [nsp]12) is a key component of the viral replication/ transcription machinery and contributes to viral genome mutation, and hence to viral adaptation. RdRp is part of a complex machinery (i.e., RdRp/nsp7/nsp8) that is fundamental to transcriptional fidelity (proofreading) (2). RdRp thus coordinates the replication of both the viral genome (which is used as the template for replication and transcription) and the shorter sgRNAs, according to the prevailing “leader-to-body” fusion model (10). Of importance, RdRp mutations at position 14,408 in European strains of SARS-CoV-2 suggest that the proofreading activity has been affected, thus altering their mutation rates (11).

Due to the dramatic global spread of COVID-19 and its associated high mortality rate, development of novel therapeutic strategies represents an unmet medical need. We therefore investigated the potential activities of long-chain inorganic polyphosphates (PolyPs) against SARS-CoV-2 viral infection.

The PolyPs are comprised of chains of a few to many hundreds of inorganic phosphates (Pi). They are ubiquitous, with prevalence in peripheral blood mononuclear cells and erythrocytes (12), and subcellularly in the nucleus, cytoplasm, plasma membranes and mitochondria (13). PolyPs are involved in of blood coagulation through activation of factor XII (14) and inhibition of the complement terminal coagulation pathway (15). Furthermore, they are involved in chelation of calcium for bone mineralisation (16) and activation of apoptosis (12). Furthermore, they act as ‘chaperone-like’ (17), and neuronal excitability molecules (18),(19),(20). Of importance, linear PolyPs have been reported to have cytoprotective and antiviral activities against human immunodeficiency virus type 1 (HIV-1) infection *in vitro* (21).

We show here that long-chain inorganic PolyPs have anti-viral activities against SARS-CoV-2 in different cellular models (i.e., Vero E6, Caco2 and human primary nasal epithelial cells). We demonstrate that intracellularly, PolyPs enhance proteasome-mediated ACE2 down-regulation, and impair transcriptional activity of RpRd during its synthesis of the major structural sgRNAs and proteins (i.e., S, E, M, N). As PolyPs are already known not to be toxic, these data open the door to clinical trials with PolyPs in humans.

## Results

### Anti-viral effects against SARS-CoV-2 of ‘preventive treatments’ with long-chain PolyPs

Anti-viral activity of linear PolyPs with average chain lengths of 15, 34, 91 Pi residues has been previously shown on HIV-infected cells (21).

To investigate potential cytoprotective effects of long-chain PolyPs, PolyP120 was used here, initially with epithelial cells derived from an African green monkey kidney (i.e., Vero E6 cells; ATCC CRL-1586), because of their high expression of ACE2 (22). The Vero cells were first incubated with 37.5 μM PolyP120, and after 30 min they were infected with SARS-CoV-2 viral particles (multiplicity of infection [MOI], 0.01) obtained from a swab of an Italian patient affected by COVID-19 (**Fig. S1A**; viral sequence shown in **Fig. S1B**). Following treatment of these cells with PolyP120, there were significantly decreased expression levels of the viral N gene (as fragments N1-3) 12 h after viral infection, compared to the vehicle control (**Fig. S1C**). Moreover, quantitative immunofluorescence of the same PolyP120-pretreated cells show reduced levels of the SARS-CoV-2 S-, E- and N-proteins (**Fig. S1D and E**). However, the RdRp protein was not down-regulated here (**Fig. S1D and E**). These data suggest that the anti-viral activity of PolyP120 can be ascribed to reduced SARS-CoV-2 entry into host cells (which is mainly mediated by ACE2), and/or inhibition of RdRp transcriptional activity.

### Anti-viral effects of PolyP120 are mediated by ACE2 down-regulation, with no cytotoxic effects

To address these hypotheses, potential binding sites for PolyP120 on both ACE2 and the viral RdRP proteins were investigated through molecular docking. These data showed the potential for binding between ACE2 and short-chain PolyPs (e.g., PolyP20), which was mostly mediated by four ACE2 amino-acid residues (Arg514, His401, His378 and Arg393; **Fig. 1A** and **Table 1**) that are conserved across different vertebrates (sequence identity, 63.6%; **Fig. 1A**).

**Table 1:** Best scores obtained through the Autodock Vina docking analysis.

| Receptor | Best score<br>(kcal/mol) |
| --- | --- |
| RdRp | -5.3 |
| ACE2 | -5.9 |
| RdRp/nsp7/nsp8 | -4.8 ± 0.2 |

**Figure 1.**
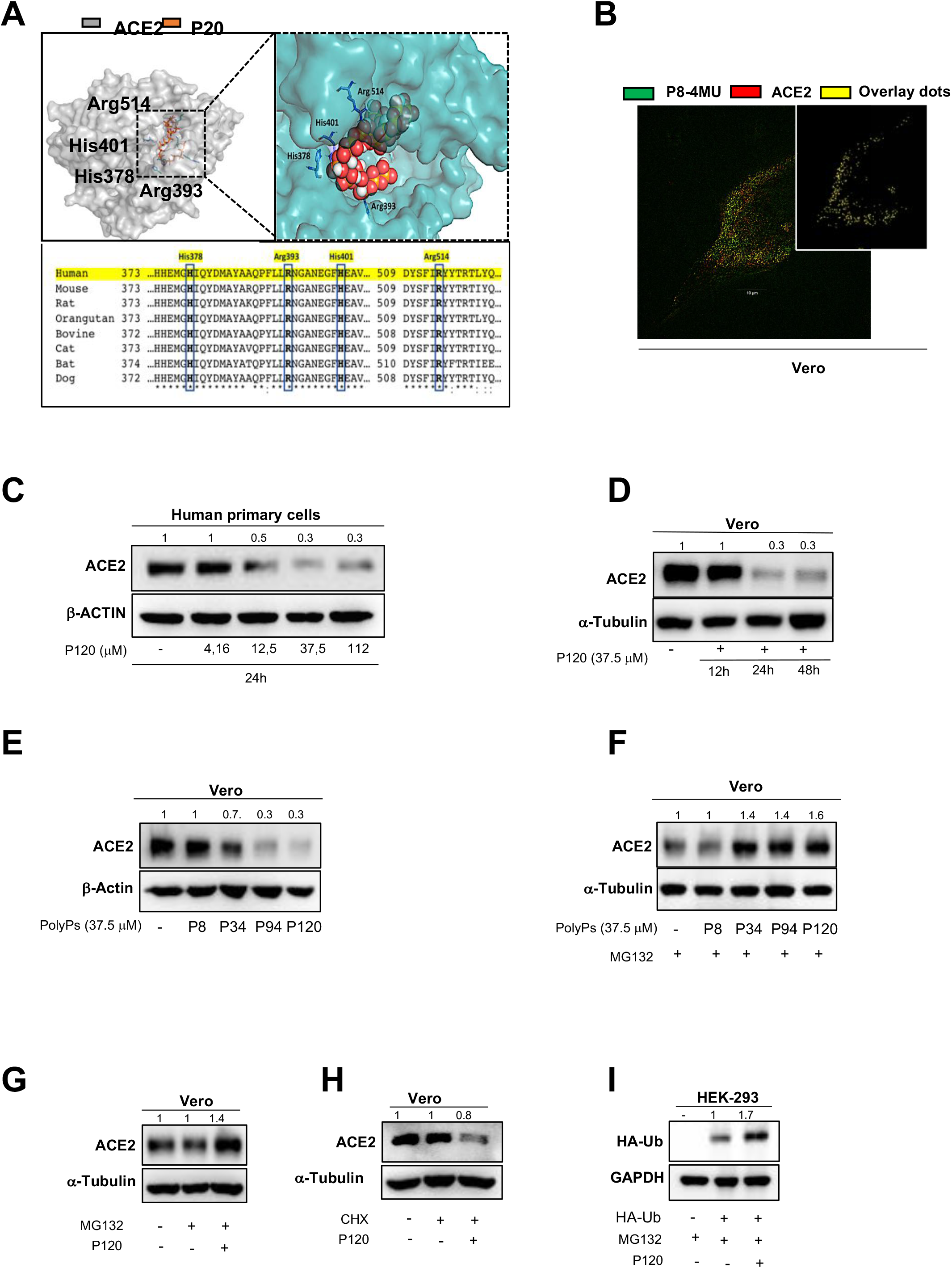
Anti-viral effects of PolyP120 are mediated by ACE2 down-regulation. **(A)** Top: Molecular docking of PolyP20 on the SARS-CoV-2 ACE2 domain (corresponding to PDB structure 6M0J chain A). ACE2 is represented as a transparent molecular surface coloured according to the electrostatic potential (left). The colour scale ranges from −10 kT/e (red) to +10kT/e (blue). The PolyP20 ligand is represented as orange sticks (left). Expanded view of the ACE2 receptor binding domain as a cyan transparent surface, to indicate the binding interface (right). Bottom: Alignment analysis of the ACE2 protein regions that contain the potential binding sites for PolyP20. The amino-acid residues mainly responsible for the interactions between ACE2 and PolyP15 are shown as blue boxes (i.e., His378, Arg393, His401, Arg514). **(B)** Vero cells treated with 37.5 μM PolyP8 conjugated with 4-methylumbelliferone (P8-4MU), and the vehicle control, were fixed and immunofluorescence was performed using an anti-ACE2 antibody. Image acquisition was made with ZEISS Elyra 7 with the optical Lattice SIM technology by using 63x oil immersion objective. The overlay spots result from colocalization analyses performed with ZEISS ZEN software (blue edition). **(C)** Human primary nasal epithelial cells were treated with different concentrations of PolyP120 (4.16, 12.5, 37.5, 112 μM), for 24 h. Vehicle-treated cells were used as negative control. Immunoblotting was performed with the antibodies indicated. **(D)** Vero cells were treated for 12 h, 24 h and 48 h with 37.5 μM PolyP120. Vehicle-treated cells were used as negative control. Immunoblotting was performed with the antibodies indicated. **(E)** Vero cells were treated for 24 h with 37.5 μM PolyP8/P34/P94/P120 (P8, P34, P94, P120). Vehicle cells were used as negative control. Immunoblotting was performed with the antibodies indicated. **(F)** Vero cells were treated for 24 h with 37.5 μM PolyP8/P34/P94/P120 (P8, P34, P94, P120) in combination with 10 μM MG132 as proteasome inhibitor. Vehicle treated cells were used as negative control. Immunoblotting was performed with the antibodies indicated. **(G)** Vero cells were treated for 24 h with 10 μM MG132 (proteasome inhibitor) and 37.5 μM PolyP120. Vehicle-treated cells with MG132 and non-treated cells were used as negative controls. Immunoblotting was with the antibodies indicated. **(H)** Vero cells were treated for 24 h with 50 μg/mL cycloheximide (protein synthesis inhibitor) and 37.5 μM PolyP120. Vehicle-treated cells with cycloheximide and non-treated cells were used as negative controls. Immunoblotting was performed with the antibodies indicated. **(I)** Plasmid containing HA-tagged Ubiquitin was transiently expressed in HEK-293 cells. After 24h from transfection cells were treated with 10 μM MG132 (proteasome inhibitor) and 37.5 μM PolyP120 for additional 24h. Vehicle-treated cells with MG132 and non-trasfected cells were used as negative controls. Immunoblotting was performed with the antibodies indicated.

The colocalisation between PolyPs and ACE2 in Vero cells was then examined. For this, the Vero cells were treated with a short-chain PolyP (i.e., PolyP8) conjugated with a fluorescent dye (i.e., 4-methylumbelliferone; MU). The immunofluorescence indicated colocalisation between PolyP8-MU and ACE2 in these treated cells (**Fig. 1B, Fig. S2A and B**). Furthermore, we investigated whether the long-chain PolyP120 colocalised with ACE2. Here, a DAPI-based approach was used to label PolyP120 (as previously described (23)), with colocalisation also seen between the intracellular PolyP120 and ACE2 (**Fig. S2C**) in these Vero cells. This was thus indicative of an interaction between PolyPs and ACE2.

As a ‘scaffold-like’ activity has been previously described for PolyPs (17), we investigated whether ACE2 protein expression can be modulated by PolyPs. Due to the high ACE2 expression levels in nasal epithelial cells (24), we used human primary cells obtained from nasal brushings of healthy subjects (**Fig. S2D**). These were treated with increasing concentrations of PolyP120. Immunoblotting demonstrated a dose-dependent reduction in ACE2 protein levels in these cells after 24 h treatments with PolyP120 (4.2-112 μM) (**Fig. 1C**). In Vero cells, the addition of 37.5 μM PolyP120 also showed a time-dependent ACE2 down-regulation (**Fig. 1D**). Of interest, the decrease in ACE2 protein levels was dependent on the number of phosphates in the PolyPs (**Fig. 1E**). Thus, to reduce the levels of ACE2 in Vero cells, they needed to be treated with PolyPs with >8 phosphate residues. Furthermore, long-chain PolyPs treatment showed greater accumulation of ACE2 upon addition of the proteasome inhibitor MG-132, compared to the vehicle controls (**Fig. 1F**), thus suggesting that the PolyPs-mediated downregulation of ACE2 occurs via proteasome. To further validate that PolyPs affect the ubiquitin–proteasome system, PolyP120 was shown not only to increase the levels of ACE2 in presence of MG-132 (**Fig. 1G**) but also to reduce its levels upon addition of the protein synthesis inhibitor cycloheximide (**Fig. 1H**). These data are further confirmed using transfection of an HA-ubiquitin plasmid construct in HEK293 cells (**Fig. 1I, Fig. S2E**).

Altogether, these data indicate that PolyP120 can bind to ACE2 and enhance its proteasome-dependent degradation, thus indicating its anti-viral mechanism of action. Of importance, the potential cytotoxicity of PolyPs were also evaluated when administered at different doses to the human primary cells, using ‘real-time’ cell proliferation assays (i.e., cell index). These data showed that only the highest dose of 112 μM PolyP120 had cytotoxic effects, as seen by decreased cell proliferation rate (**Fig. S2F**). However, Caspase 3 enzymatic assays did not show activation of the apoptotic cascade by PolyP120 (**Fig. S2G**), thus further confirming the lack of toxic effects of PolyP120.

### PolyP120 inhibits transcription of viral sgRNAs through inhibition of RdRp

To gain further insight into the anti-viral mechanisms of PolyPs in their inhibition of viral gene transcription, we investigated potential interactions with RdRp. Molecular docking analysis showed putative binding sites for PolyP15 on the RdRp protein **(Fig. S3A** and **Table 1)**. Further docking studies performed with RdRp in its active replicating form (i.e., RdRp/nsp7/nsp8 complex; **Table 1**) illustrated possible binding modes with PolyP20. These are mediated through seven RdRp amino-acid residues located within the RNA-binding site of RdRp (K4892, K4937, R4945, D5152, R5228, D5237, and K5241; **Fig. 2A**) that are conserved in RdRp of other viruses in the *Coronoviridae* family (e.g., SARS-CoV, MERS-CoV; **Fig. 2A)**, and potentially in the main families of positive-sense single-stranded RNA viruses from other hosts (e.g., *Nidovirales*; as listed at *https://www.ncbi.nlm.nih.gov/genomes/GenomesGroup.cgi?taxid=76804*).

**Figure 2.**
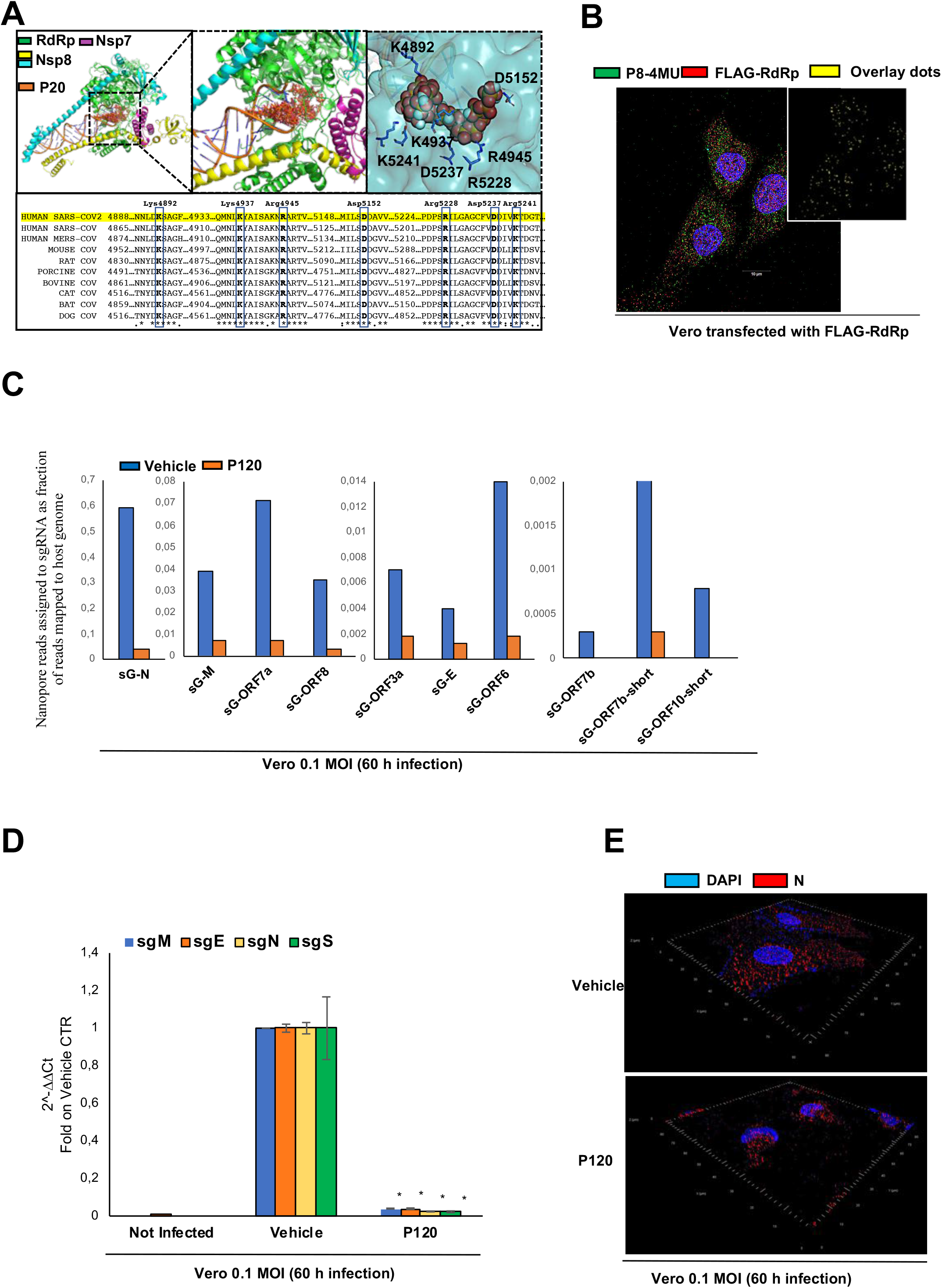
PolyP120 inhibits transcription of viral sgRNAs through inhibition of RdRp. **(A)** Top: Molecular docking of PolyP20 (P20) on SARS-CoV-2 RdRp (corresponding to PDB structure 6M71) (left). RdRp is represented as a molecular surface coloured according to electrostatic potential. Colour scale ranges from −10 kT/e (red) to +10kT/e (blue). The ligand PolyP20 is shown as red balls. Expanded view of molecular docking is shown (middle panel), with further expanded view (right) of RdRp as a cyan transparent surface, to indicate the binding interface. Bottom: Alignment analysis of the RdRp protein (nsp12) region that contains the potential binding sites with PolyP20. The amino-acid residues mainly responsible for interactions between RdRp and PolyP20 are shown in blue boxes (i.e., Lys4937, Arg4945, Asp5152, Arg5228, Asp5237, Arg5241). **(B)** Vero cells transfected with FLAG-RdRp and treated with 37.5 μM PolyP8 conjugated with 4-methylumbelliferone (P8-4MU), and the vehicle control, were fixed and immunofluorescence was performed using an anti-RdRp antibody. Image acquisition was made with ZEISS Elyra 7 with the optical Lattice SIM technology by using 63x oil immersion objective. The overlay spots result from colocalization analyses performed with ZEISS ZEN software (blue edition). **(C)** Quantification of direct RNA sequencing reads (through Nanopore technology) assigned to sgRNAs expressed as fractions of the reads mapped to the host genome. Note the different scales for the Y axes for each sgRNA. **(D)** Vero cells were infected with SARS-CoV-2 viral particles (0.1 MOI), with non-infected cells as negative control of infection. After 24 h, cells were treated with 37.5 μM PolyP120 or vehicle, and 36 h later they were lysed and RNA was extracted. Ct values from real-time RT-PCR are shown for expression of subgenomic (sg) viral RNAs, as indicated. **(E)** Immunofluorescence with antibodies against N protein on Vero cells infected with SARS-CoV-2 viral particles (0.1 MOI) and treated with PolyP120. The SIM image was acquired with Elyra 7 and 3-dimentional (3D) reconstruction of Z stacking data were performed by using ZEISS ZEN software (blue edition). Magnification 63×.

Using immunofluorescence, colocalisation between PolyP8-MU and RdRP was indicated for Vero cells transfected with a FLAG-RdRp plasmid construct (**Fig. 1B** and **Fig. S3B and C, on left panel**). Moreover, immunofluorescence of SARS-CoV-2-infected Vero cells showed colocalisation between intracellular PolyP120 and RdRp protein (**Fig. S3C on right panel**), thus indicating the potential for interactions.

Whether the downregulation of the viral genes mediated by PolyPs was due to inhibition of the viral transcription process was also investigated. Thus, we used total RNA samples from oropharyngeal swabs of eight Italian patients with COVID-19 (**Table 2**) and examined modulation of amplification of the viral N1-3 gene fragments in the presence of PolyP120 (**Fig. S3D**). These data showed increase in Ct values for both N1 and N3, and a total inhibition of the N2 gene fragment (i.e., undetermined Ct) when the reactions were performed in presence of 37.5 μM PolyP120 (**Fig. S3E**). Of interest, addition of PolyP120 did not alter amplification of the N gene fragments when external S RNA controls (N1/2/3-O_2_-methyl) were used (**Fig. S3F**). These *in-vitro* data indicated that PolyP120 requires the host proteins and/or carriers for inhibition of viral gene transcription.

**Table 2.** Clinical information and Ct values for the cohort of Italian patients with COVID-19. All of the samples were from oropharyngeal swabs that originated from Naples, Italy.

| Sample code | Patient | | Sample arrival<br>date<br>(dd/mm/yyyy) | Target | Vehicle<br>(Ct) | 37.5 μM<br>PolyP120<br>(Ct) |
| --- | --- | --- | --- | --- | --- | --- |
|  | Age<br>(years) | Sex |  |  |  |  |
| 20200455403 | 34 | Female | 22/04/2020 | N1 | 20.68 | 24.46 |
|  |  |  |  | N2 | 20.16 | n.a. |
|  |  |  |  | N3 | 21.66 | 23.23 |
| 20200405430 | 64 | Male | 20/04/2020 | N1 | 32.01 | 32.26 |
|  |  |  |  | N2 | 32.09 | n.a. |
|  |  |  |  | N3 | 32.28 | 33.68 |
| 20200406267 | 65 | Male | 20/04/2020 | N1 | 32.07 | 34.57 |
|  |  |  |  | N2 | 32.50 | n.a. |
|  |  |  |  | N3 | 31.61 | 35.49 |
| 20200405509 | 73 | Female | 17/04/2020 | N1 | 32.18 | 34.21 |
|  |  |  |  | N2 | 32.41 | n.a. |
|  |  |  |  | N3 | 33.01 | 33.27 |
| 20200406280 | 59 | Male | 20/04/2020 | N1 | 30.32 | 32.97 |
|  |  |  |  | N2 | 30.54 | n.a. |
|  |  |  |  | N3 | 30.26 | 34.63 |
| 20200405519 | 42 | Male | 22/04/2020 | N1 | 32.60 | 33.77 |
|  |  |  |  | N2 | 33.43 | n.a. |
|  |  |  |  | N3 | 33.15 | 35.08 |
| 20200405527 | 76 | Male | 17/04/2020 | N2 | 31.77 | 33.04 |
|  |  |  |  | N2 | 32.21 | n.a. |
|  |  |  |  | N3 | 32.08 | 33.87 |
| 20200406213 | 56 | Male | 20/04/2020 | N1 | 32.11 | 36.36 |
|  |  |  |  | N2 | 33.02 | n.a. |
|  |  |  |  | N3 | 32.24 | 35.82 |

n.a., not amplified

Then, to further dissect out transcriptional inhibition mediated by PolyP120 on RdRp, we focused on its discontinuous transcription, taking into account four structural sgRNAs (i.e., sgS, sgE, sgM, sgN). For this purpose, SARS-CoV-2–infected Vero cells were treated with PolyP120 and then analysed for transcription of these sgRNAs (**Fig. S4A**). These data confirmed a reduction in the N gene fragments, RdRp and S transcripts in PolyP120-treated cells (**Fig. S4B and C**). Additional data were obtained using Nanopore technology and direct RNA sequencing of the RNAs extracted from viral infection of these Vero cells showing decreased levels of the sgRNAs (**Fig. 2C** and **Table S2**; see Methods and CEINGE portal). Furthermore, using RT-PCR with SYBR-Green, we also confirmed these data showing decreased levels of sgS, sgE, sgM and sgN (**Fig. 2D**). Moreover, immunoblotting performed with PolyP120-treated SARS-CoV-2-infected Vero cells also showed dramatic reductions in both ACE2 and N-protein levels (**Fig. S4D**).

This ‘therapeutic’ anti-viral action of PolyP120 was also confirmed by immunofluorescence, which showed significant reduction of ACE2 and the SARS-CoV-2 proteins (i.e., N, E, S) in these PolyP120-treated Vero cells (**Fig. 2E, Fig. S4E and F,** and **Fig. S5**). Furthermore, through Hulu-FISH in situ hybridization approach combined to quantitative Immunofluorescence we also detect a reduction of gRNA-Spike and ACE2 protein in SARS-CoV-2-infected Vero cells upon addition of PolyP120 (see **Fig. 4A**). Altogether, these data show the mechanisms of action of the PolyPs through reduction in the transcriptional activity of RdRp, which appears to be due to their binding to its active enzymatic core (see **Fig. 2A**).

**Figure 4.**
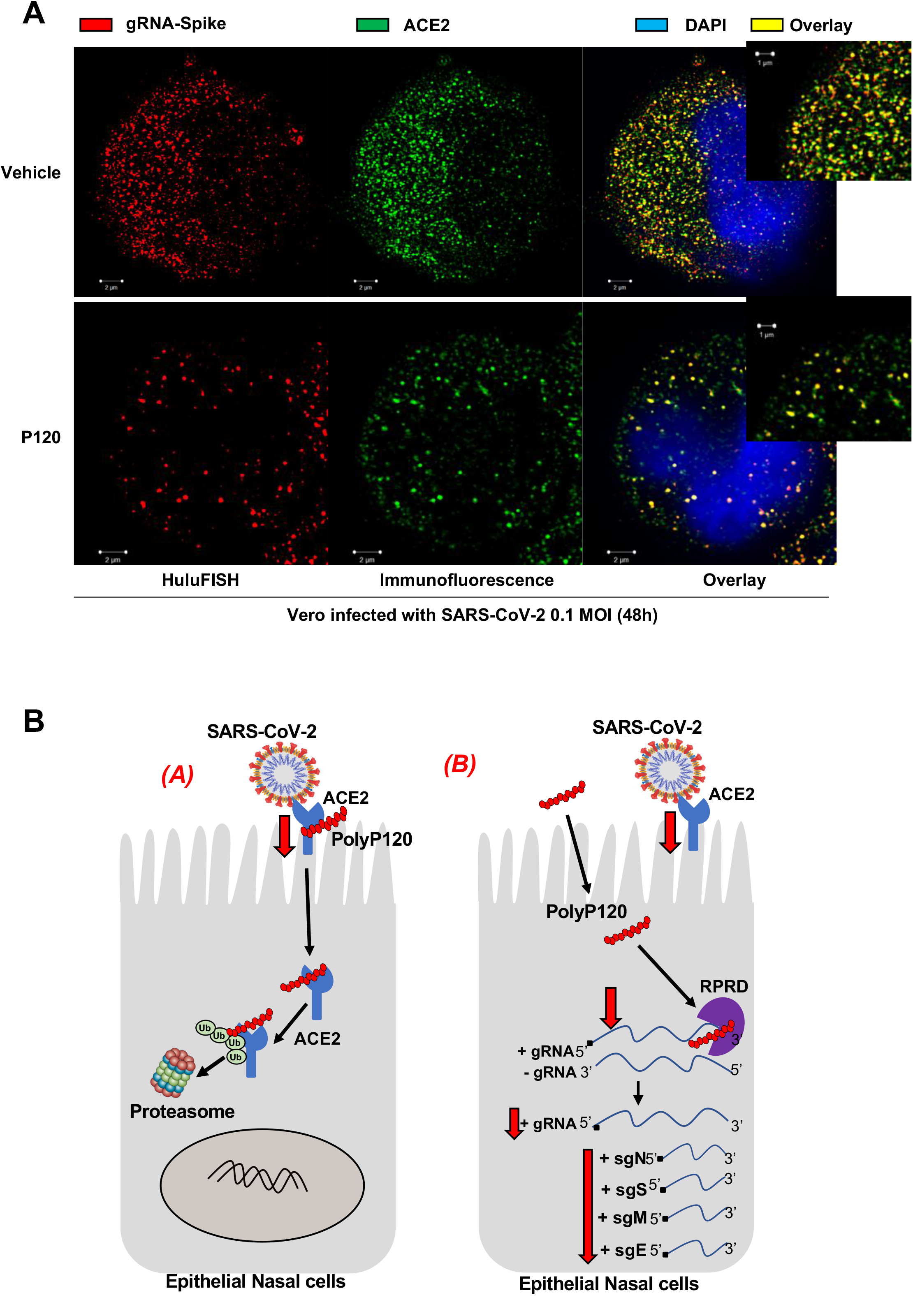
Intracellular mechanism of action of PolyP120. **(A)** HuluFISH with Pan-SARS-CoV-2 probe (red) coupled to Immunofluorescence staining with antibody against ACE2 protein (green) on Vero cells infected with SARS-CoV-2 (Genbank: MT682732.1; MT890669) particles (0.1 MOI) and treated with 37.5 μM PolyP120. The data shown a substantial *in cell* downregulation of gRNA-Spike (red) and ACE protein (green) expression in Poly120-treated versus vehicle cells. The SIM image was acquired with Elyra 7 and processed by using ZEISS ZEN software (blue edition). Magnification 63×. **(B)** Cartoon representation to illustrate our hypothesis for the anti-viral actions of PolyPs (e.g., P120). PolyPs can act via proteasome-mediated degradation of ACE2 *(A)*, and can impair RpRd transcription of sgRNA genes viral transcription *(B)*. Thus PolyPs can act extracellularly via direct interactions with the host-cell ACE2, as the extracellular receptor for interaction with the Spike of the virus, or intracellularly through its interactions with RdRp.

Of interest, we also noted down-regulation of ACE2 at the transcriptional level (**Fig. S4C**). Treatment with the transcriptional inhibitor actinomycin D showed significant decrease in ACE2 mRNA expression levels in the Vero cells after 24 h (**Fig. S4G**), thus also suggesting transcriptional inhibition mediated by PolyPs.

### Therapeutic treatments with long-chain PolyPs

To investigate the potential ‘therapeutic’ effectiveness of PolyPs, additional studies were performed in which PolyPs at different chain length were added after SARS-CoV-2 infection. For this purpose, SARS-CoV-2-infected Vero cells were incubated with PolyPs (starting from the 120 phosphate residues of PolyP120, and including PolyP126 and PolyP189), and then analysed for expression levels of the viral N gene (**Fig. S6A**). No significant differences were seen between these different chain length PolyPs for the decreased expression of the viral N gene (N1-3), compared to the vehicle control (**Fig. S6B**).

Furthermore, similar data were obtained for Vero cells infected with SARS-CoV-2 particles obtained from a Korean patient (EPI_ISL_407193; **Fig. S1B**), by measuring viral RdRp and E genes expression upon addition of PolyP120 at different concentrations (9.38, 18.75, 37.50 μM) (**Fig. S6C and D**). The increased Ct values for the RdRp gene in these Vero cells treated with PolyP120 indicated inhibition of SARS-CoV-2 infection, including for the lowest PolyP120 concentration used (**Fig. S6D)**.

As different mutations have been identified in Europe for the RdRp, S and ORF1ab genes (11), we also performed sequencing analysis focusing on the viral regions that flank these hot-spot mutation sites between the Italian and Korean SARS-CoV-2. These data showed non-synonymous variants in these genes (i.e., RdRp, S, ORF1ab) in the viral genome from an Italian patient compared to that from the Korean patient (**Fig. S2A, red arrows**). Furthermore, these variants were conserved across Italian viral sequences, compared to Asian viral sequences (**Fig. S2A)**, as previously described (25). Importantly, the mutation in RdRp (at position 14408) was associated with an increased mutation rate, which would appear to be due to an alteration in the proofreading that is finely regulated by the RdRp/nsp7/nsp8 machinery complex (11). Our data show that PolyP120 can decrease the expression of these viral genes in the infected Vero cells also in the presence of this SARS-CoV-2 point mutation. In this regard, our docking analysis suggested that the mutated site in RdRp falls outside of its RNA-dependent transcriptional activity domain and its potential binding sites with the PolyPs (**Fig. S6E**). This thus supports the hypothesis that the PolyPs impair the RNA-dependent transcriptional activity rather than the proofreading activity.

### Anti-viral actions of PolyP120 in human cell lines decrease the COVID-19 cytokines storm

To determine further whether increased PolyPs size, in terms of Pi residues, has any effects on their antiviral activity, human primary cells were pretreated with PolyP120 and PolyP126, then infected with SARS-CoV-2 at the different viral loads of 0.01 and 0.16 MOI, and analysed for expression levels of the viral N gene (**Fig. S7A**). These data indicated that PolyP126 shows similar anti-viral activity to PolyP120 in these human cells infected with SARS-CoV-2 (**Fig. S7B**). Altogether, these data suggest that following the pretreatment of human primary nasal epithelial cells, the longer-chain PolyPs (e.g., PolyP120, PolyP126) have anti-viral effects through inhibition of N gene transcription. This preventive antiviral action was also confirmed in SARS-CoV-2–infected human colorectal carcinoma cells (i.e., Caco2 cells), where PolyP120 pre-treatment showed a decrease for the viral genes (N, RdRp) (**Fig. S7C to E**) and ACE2 protein (**Fig. S7F**). These data thus suggest that the mechanism of action of PolyP120 is conserved across different human cellular models.

Furthermore, we also tested the ‘therapeutic’ effectiveness of PolyP120 in SARS-CoV-2-infected human primary nasal epithelial cells (**Fig. 3A, Fig. S8A**). The PolyP120 treatment showed decreased levels of the N1-3 gene fragments (**Fig. S8B**), and reductions in the other viral genes (i.e., RdRp, S, E) and ACE2 (**Fig. S8C**). Furthermore, immunoblotting confirmed total inhibition of the ACE2 and N protein levels in these treated human primary nasal epithelial cells (**Fig. 3B**).

**Figure 3.**
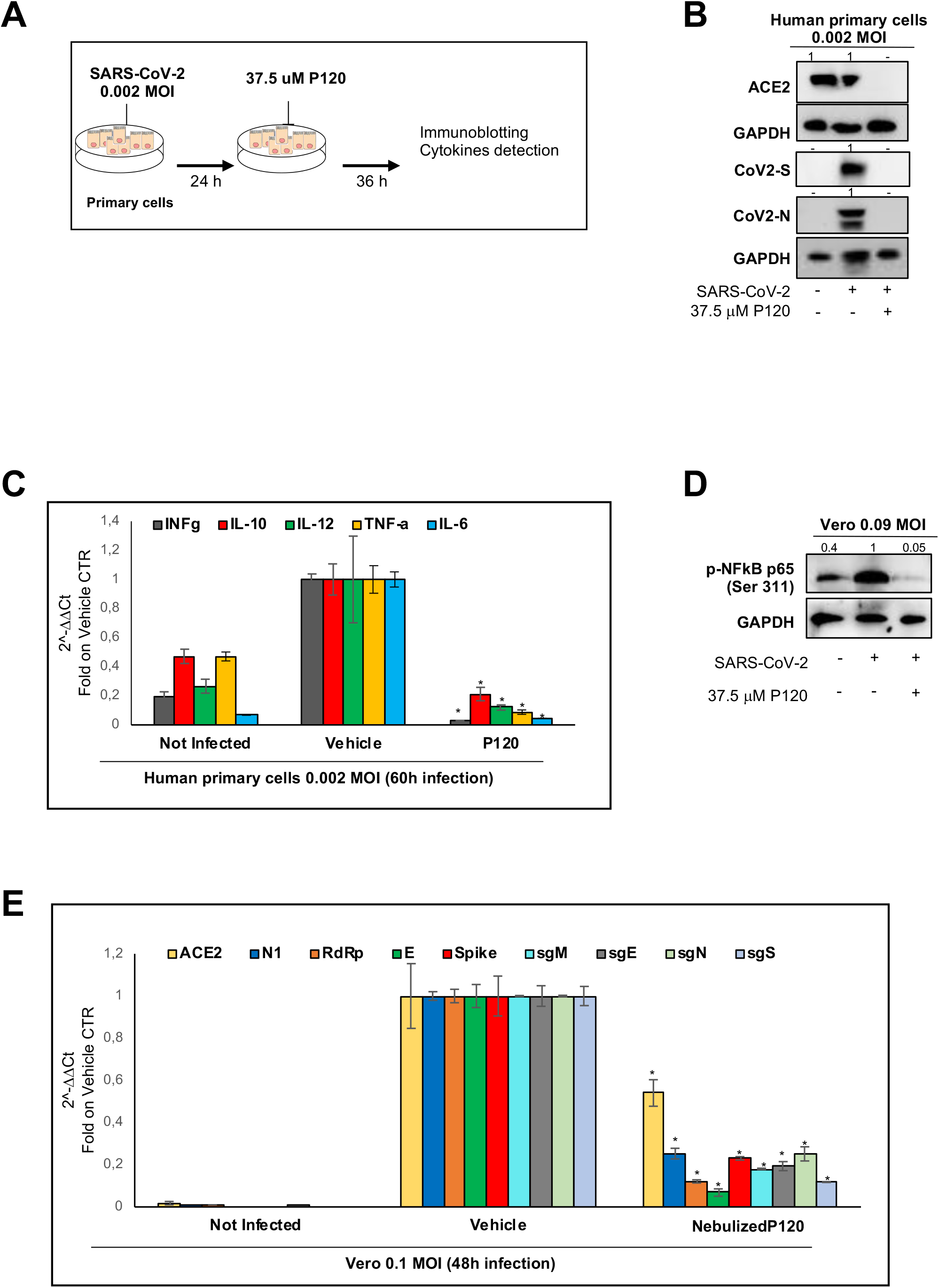
Therapeutic efficacy of PolyP120 delivered through a nonambulatory nebuliser system. **(A)** Experimental plan. Human primary nasal epithelial cells (425,000) were infected with SARS-CoV-2 viral particles (0.002 MOI), with non-infected cells as negative control of infection. After 24 h, cells were treated with 37.5 μM PolyP120, and 36 h later they were lysed and RNA was extracted. **(B)** Immunoblotting of non-infected and SARS-CoV-2-infected human primary nasal epithelial cells treated with 37.5 μM PolyP120 and vehicle, using antibodies as indicated. **(C)** Ct values from real-time RT-PCR following RNA extraction, showing expression of interferon-γ, TNF-α and interleukins IL-6, IL-12, IL-10. **(D)** Immunoblotting of SARS-CoV-2-infected Vero cells treated with 37.5 μM PolyP120 and vehicle, using antibodies as indicated. **(E)** Vero cells were treated with nebulised 37.5 μM PolyP120. After 1 h, these cells were infected with SARS-CoV-2 viral particles (0.14 MOI), with non-infected cells as negative control of infection. After 48 h, the cells were lysed and RNA was extracted. Ct values from real-time RT-PCR are shown for expression of the ACE2, CoV-2 N1, E, RdRp, S genes, and subgenomic (sg) viral RNAs, as indicated.

Patients with COVID-19 also show increased circulating levels of inflammatory cytokines, including interferon-γ and interleukin (IL)-6, IL-10 and IL-12 (26). As the PolyPs have been previously described as having anti-inflammatory functions (14), we investigated the potential modulation by PolyP120 of these cytokines in human primary cells infected with SARS-CoV-2. Indeed, real-time PCR analyses showed significant reductions in interferon-γ, IL-6, IL-10 IL-12 and tumour necrosis factor-α (**Fig. 3c**). The anti-inflammatory action exerted by PolyPs could be ascribed to the inhibition of NF-kB pathway, as shown by reduced levels of phosphorylated (Ser311)-p65 in SARS-CoV-2-infected Vero cells after treatment with PolyP120 (**Fig. 3D**).

As the PolyPs have been previously reported to be associated with the inflammatory pathways that are mainly regulated by NF-κB (27) and mTOR (28), we hypothesise that this down-regulation of the cytokines can be ascribed to modulation of the inflammatory cascades in an autocrine and paracrine manner by PolyP120.

### Delivery of PolyP120 through a nonambulatory nebuliser system

Due to the respiratory failure experienced by patients with COVID-19, new drug formulations that are appropriate for aerosol inhalation, and thus more direct lung delivery, represent an unmet medical need. For this reason, we tested the delivery of PolyP120 into Vero cells via a nebuliser system. Here, 37.5 μM PolyP120 was nebulised for 2 min (condensation rate, 1 mL/min) into the tissue culture flask containing the attached Vero cells in the absence of medium. After nebulising, medium was added back to the cells, and the flasks were left for 12 h under growth conditions (see **Video 1, Supplemental Material**). Entrance of the PolyP120 into the cells was confirmed by immunofluorescence using DAPI, thus showing the efficacy of this delivery for the nebulised PolyP120 into the cells (**Fig. S8D**).

We then investigated whether this PolyP120 aerosol formulation maintains the antiviral activity of PolyP120 on Vero cells. Thus, the nebulised PolyP120 was similarly applied to Vero cells that were then infected with SARS-CoV-2 viral particles (**Fig. S8E**). Real-time PCR analysis indeed showed down-regulation of the N gene fragments and reduced levels of ACE 2 and the viral genes, and all of the structural sgRNAs (**Fig. 3E**). This thus indicated that PolyP120 delivered in this aerosol formulation maintained its anti-viral action against SARS-CoV-2 infection.

## Conclusions

Here, we have shown that long chain length PolyPs (i.e., PolyP120, with 120 phosphate residues) can inhibit the transcription of SARS-CoV-2 viral genes (**Fig. 4A**) impairing the replication of the virus in the host cell. Future diagnostic use of this technology *in vivo* will be of great impact monitoring the disease status.

Of importance, we have described two intracellular mechanisms of action behind this anti-viral function of PolyPs, as summarised in **Fig. 4B**. The first mechanism involves time- and dose-dependent proteasome-mediated degradation of ACE2 protein in the presence of long chain length PolyPs. This appears to be due to the binding of the PolyPs to ACE2, which should be mainly mediated by four amino-acid residues of ACE2 that are also conserved across different species: Arg514, His 401, His 378 and Arg393. This is of importance to avoid further spillover. We also note the role of PolyPs as a source of Pi to increase the pool of ATP available to enhance the ubiquitin–proteasome system (29).

Interestingly, we have also shown that the SARS-CoV-2-infected cells have reduced ACE mRNA expression levels after treatment with PolyP120, we think this is because the expression of ACE2 is regulated by the HNF1α transcriptional factor (30) and epigenetically by SIRT1 (31). These phenomena need further investigations.

Of note, in directing any treatment here at ACE2 as the receptor for SARS-CoV-2 entry into cells, the use of ACE inhibitors has been discouraged for patients with COVID-19. This is because of the consequent increased levels of ACE2 mRNA, which might facilitate engagement and entry of SARS-CoV-2 into cells (32). However, the future use of PolyPs might indeed overcome this side effect of the use of ACE inhibitors. Thus, the combination of ACE inhibitors with PolyPs can be postulated for the treatment of patients with COVID-19.

The second mechanism of action presented here involves transcriptional inhibition of RdRP (**Fig. 4B)**. The PolyPs appear to bind to viral RdRp from SARS-CoV-2, mainly through seven of its amino-acid residues located within the RdRp RNA-binding site. PolyP120 appears to decrease the expression of the main SARS-CoV-2 viral genes through this mechanism. Of importance, the transcription of the main viral sgRNAs is also impaired confirming their inhibitory actions on RdRp transcription. This inhibition also occurs in the presence of non-synonymous variants (at position 14408) that have been shown to affect the integrity of the proofreading activity, which further supports the importance of PolyPs to universally impair viral gene transcription. The amino-acid residues in RdRp that are potentially involved in the binding with PolyPs have also been conserved through evolution in different species, and thus they might also be conserved in the other positive-sense singlestranded RNA viruses from other hosts (e.g., *Nidovirales;* as listed at *
https://www.ncbi.nlm.nih.gov/genomes/GenomesGroup.cgi?taxid=76804*). These findings open the way for the potential use of PolyPs against other viral infections.

The various concentrations of PolyP120 tested here are also shown not to have any toxic effects, as seen by monitoring of the caspase 3 enzymatic activation in human primary nasal epithelial cells. However, the highest concentration of PolyP120 tested (i.e., 112 μM) blocked proliferation of these primary cells. This anti-proliferative effect might be due to the inhibition of the MAP-kinase cascade (via ERK), which has previously been reported for inorganic PolyPs (33).

PolyPs have also been shown to be key modulators of inflammatory pathways, with actions mainly via the NF-κB cascade (27). Here, we have shown that PolyP120 decreases the levels of the main cytokines (i.e., interferon-γ, IL-6, IL-10, IL-12) that are part of the cytokine storm often seen for patients with COVID-19 (26) by affecting NF-kB cascade. Among these cytokines, IL-6 levels in patients with COVID-19 are significantly elevated, and are associated with adverse clinical outcomes. While inhibition of IL-6 levels with tocilizumab appears to be effective and safe in preliminary investigations, results need to be evaluated of ongoing clinical trials to better define the role of tocilizumab in COVID-19 and its routine clinical application (34).

Pre-existing T-cell immunity to SARS-CoV-2 might be relevant also, because it might influence COVID-19 disease severity. It is possible that people with high levels of preexisting memory CD4+ T-cells that recognize SARS-CoV-2 can mount a faster and stronger immune response upon exposure to SARS-CoV-2, and thereby limit the disease severity.

Furthermore, in COVID-19 infections, microcapillary clotting in the lung generates acute pulmonary embolisms, which impairs O2 acquisition by the lungs, thus resulting in insufficient respiratory function. Of importance, low molecular weight heparin sulphate given to the patients reduces their pain, improves their clinical status, reduces their IL-6 release, and stabilises their oxygen exchange (35). At this time it is not clear if PolyP can suppress the complement protein via the terminal pathway (15) and enhance the clotting formation. For these reasons, cautions should be taken to treat late stage disease patients with PolyP.

Comorbidity, asymptomatic and symptomatic patients can use different immune systems to fight SAR-CoV-2, where memory CD4+ T-cells have been shown to have a role (36). The use of PolyPs can enhance the immunological memory, including T-cell activation, and this can be monitored *in vivo* in further clinical trials. These will be issues for future efforts.

Patients with COVID-19 suffer from respiratory failure. So new drugs formulated as aerosols should represent a promising therapeutic approach. Interestingly, we also show here that the efficacy of PolyPs is maintained when they are nebulised for their delivery to the cells, as the PolyP120 still showed antiviral activity against SARS-CoV-2 in infected Vero cells—Thus, we envision the aerosol delivery of PolyP120, for the treatment of patients with respiratory failure. The final formulation will be discussed elsewhere.

Finally, these data show ‘preventive’ effects of the long chain length PolyPs against SARS-CoV-2 infection. In this regard, the panel on Food Additives and Flavourings has already provided information about the safety of phosphates as food additives (i.e., E450–452) (37).

To date, among the drugs tested on COVID-19 patients, the safety and antiviral activity of Remdesivir (GS-5734) is emerging in two completed phase 3 clinical trials (NCT04292899, NCT04292730). Assuming no toxicity of PolyP120, we hypothesize its rapid use in clinics in early phase of COVID-19 disease (2) alone or in combination with Remdesivir. Thus, the use of PolyP120 against infections with SARS-CoV-2 or other new expected virus can be envisioned for the near future.

### Ethic Committee approval

Ethical Committee approvals for the Covid19 samples use in this study:

i. Protocol n. 141/20, date 10/04/2020, TaskForce CEINGE TaskForce Covid19; Azienda Ospedaliera Universitaria Federico II, Direzione Sanitaria, protocol n. 000576 of 10/04/2020.
ii. Protocol n. 157/20, date 22/04/2020, GENECOVID, the experimental procedures within the use of SAR-CoV-2 in BLS3 laboratory are authorized by Ministero della Sanità and Dipartimento Di Medicina Molecolare e Biotecnologie Mediche, Università degli Studi di Napoli Federico II and Azienda Ospedaliera Universitaria Federico II, Direzione Sanitaria protocol n. 0007133 of 08/05/2020.
iii. Protocol n. 18/20, date 10/06/2020, Genetics TaskForce CEINGE TaskForce Covid19; Azienda Ospedaliera Universitaria Federico II, Direzione Sanitaria protocol n. 000576 of 10/04/2020.

## Methods

### Cell culture

Vero cells were kept in DMEM (41966-029; Gibco) with 10% FBS (10270-106; Gibco), 2 mM L-glutamine (25030-024; Gibco), and 1% penicillin/streptomycin (P0781; Sigma). Freshly isolated human nasal epithelial cells were collected by nasal brushing of healthy donors (as previously described (38)). These were cultured in PneumaCult (#05009; StemCell Technologies) with 2 mM L-glutamine (25030-024; Gibco), and 1% penicillin/streptomycin (P0781; Sigma). Caco2 cells were cultured in EMEM (M2279, Sigma-Aldrich) with 10% FBS (10270-106; Gibco), 2 mM L-glutamine (25030-024; Gibco), and 1% penicillin/streptomycin (P0781; Sigma).

For proteasome inhibition treatment Vero E6 cells were incubated in DMEM (10% FBS) containing 10uM Mg132 (M8699; Sigma-Aldrich) for 24h; for protein synthesis inhibitor treatment, Vero E6 cells were incubated in DMEM (10% FBS) containig 50 ug/mL cycloheximide (C7698; Sigma-Aldrich) for 24h. For transcription inhibition treatment Vero E6 cells were incubated in DMEM (10% FBS) containing 10 μg/mL Actinomycin D (A1410; Sigma-Aldrich) for 24h. Following each treatment, cells were analyzed by Western Blotting as described below.

HEK-293T or Human primary nasal epithelial cells were transiently transfected with the plasmid DNA constructs (i.e. HA-ubiquitin plasmid, pGBW-m4134165 RdRp Amp^r^ plasmid) using HilyMaxTransfection Reagent (H357-10; Dojindo) according to the manufacturer instructions. Hek 293T transfected cells were then treated for 24h with Polyp120 and harvested for Western blotting 48 h after transfection. Human primary nasal epithelial cells were then treated for 12h with PolyP8-MU and harvested for Immunostaining 48 h after transfection.

### In-vitro treatment and infection

#### Preventive treatment

Vero cells or human primary nasal epithelial cells were plated in T25 flasks (6 ×10^5^ cells/flask) for treatment with 37.5 μM PolyPs (from P94 to P126). After 20 min, these pretreated cells were infected with SarsCov2 viral particles obtained from a frozen swab from an Italian patient positive for COVID-19. Non-infected cells were used as the negative control. After 12 h of infection, the cells were lysed and their RNA was extracted. Vehicle-treated cells were used as the negative control for this treatment. These experiments were performed in a B3 authorised laboratory.

#### Therapeutic treatment

Vero cells (4 × 10^5^) were plated in six-well plates for the viral infection with SarsCov2 viral particles, which were obtained from a frozen swab from Italian or Korean patients positive for COVID-19. Non-infected cells were used as the negative infection control. After 24 h, the infected cells were treated with PolyPs of different chain lengths (up to PolyP189) or PolyP120 at different concentrations (9.38, 18.75, 37.5 μM). Vehicle-treated cells were used as the negative treatment control. After 24 h of PolyPs treatments (i.e., 48 h after infection), the Vero cells were lysed and their RNA was extracted. These experiments were performed in a BLS3 authorised laboratory.

#### Nebulised PolyP120 treatment

Vero cells (6 × 10^5^) were plated in a T25 flask. The PolyP120 (37.5 μM) was dissolved in 2 mL water solution and nebulised for 2 min using a nebuliser system (mesh nebulizer; YM – 252; input: 5 V/1 A; oscillation frequency, 110 kHz; atomised particles, <5 μM; condensation rate, 1 mL/min). Following removal of the medium from the flask, the nebulised PolyP120 was directed into the flask. The medium was then replaced after the nebulising of the PolyP120. PolyP120 in the cells was detected by immunofluorescence using DAPI. These Vero cells treated with nebulised PolyP120 were then infected with the SARS-CoV-2 viral particles.

### Real-time RT assays

#### N1-3 detection

RNA samples were extracted with TRIzol RNA Isolation Reagent (#15596018; Ambion, Thermo Fisher Scientific), according to the manufacturer instructions. Real-time RT-PCR was performed using ‘quanty COVID-19’ kits (Ref. RT-25; Clonit; US Food and Drug Administration ‘*in-vitro* diagnostic’ (IVD) approved). These kits allow specific quantitative detection of the N1, N2 and N3 fragments (from the Sars-CoV-2 N gene), using differentially labelled target probes. These runs were performed on a PCR machine (CFX96; BioRad) under the following conditions: 25 °C for 2 min; 50 °C for 15 min; 95 °C for 2 min; 95 °C for 3 s; 55 °C for 30 s (×45 cycles).

#### RdRp and E detection

RNA samples were extracted with TRIzol RNA Isolation Reagent (#15596018; Ambion, Thermo Fisher Scientific), according to the manufacturer instructions. Real-time RT-PCR was performed using ‘RealQquality RQ-2019-nCoV’ kits (Ab Analitica; IVD approved) for detection of the RdRp and E genes (from the SarsCov2 N gene). These runs were performed on a PCR machine (CFX96; BioRad) under the following conditions: 48 °C for 10 min; 95 °C for 10 min; 95 °C for 15 s; 60 °C for 1 min (×45 cycles).

#### ACE2, N1, RdRp, E, S, sgM, sgE, sgN, sgS detection

RNA samples were extracted with TRIzol RNA Isolation Reagent (#15596018; Ambion, Thermo Fisher Scientific), according to the manufacturer instructions. Reverse transcription was carried out using ‘5× All-in-one RT Mastermix’ (#g486; ABM), following the manufacturer instructions. The cDNA preparation was through the cycling method, as follows: incubation of the complete reaction mix at 25 °C for 5 min; at 42 °C for 30 min; at 85 °C for 5 min; and hold at 4 °C. The reverse transcription products (cDNA) were amplified by quantitative real-time PCR using a real-time PCR system (7900; Applied Biosystems, Foster City, CA, USA). The relative expression of the target genes was determined using the 2-ΔCt method. All of the data are presented as means ±standard error of two to three replicates. The target genes were detected using a Brightgreen 2× qPCR Mastermix low-rox (#Mastermix-lr; ABM.). The details of the primers used in these assays are given in Supplementary Table 1 in the Supplementary Information.

#### Nanopore RNA direct sequencing

To carry out the direct sequencing of the RNA with nanopore technology, we prepared the RNA library using direct RNA sequencing kits (SQK-RNA002; Oxford Nanopore kits), following the manufacturer instructions. Briefly, we used poly(dT) adapter and SuperScript II Reverse Transcriptase (Thermo Fisher Scientific) to generate RNA-DNA hybrids. Agencourt RNAClean XP magnetic beads (BECKMAN) were used to purify the RNA-DNA duplexes. Subsequently, RNA-DNA hybrids were ligated to Nanopore sequencing adapters (RMX) using T4 DNA ligase (NEB E6056) prior to sequencing. After another step of purification using Agencourt RNAClean XP magnetic beads, the library concentration was estimated with Qubit assay (Thermo Fisher). Then, the Nanopore direct RNA library was loaded into the flow cell (R9.06) using MinION for 26 h. The motor protein RMX pulls the 3’-end of the RNA strand into the Nanopore channel (*39*). Changes in the ionic current are then detected at each pore by a sensor.

#### Classifying ONT reads from subgenomic RNAs

Reads from Nanopore direct sequencing were initially identified by alignment to a set of reference sequences using Minimap2 version 2.17 (40), with the ‘long-read spliced alignment’ preset. The reference sequences include SARS-CoV-2 viral genome (GenBank, NC_045512), the *Chlorocebus sabaeus* host genome (assembly ChlSab 1.1), and the human ribosomal DNA repeat (GenBank, U13369.1). Reads identified as viral were then realigned to the same viral genome sequence using Minimap2, with the following options: -k 8 -w 1 -g 30000 --max-chain-skip 40 -p 0.7 -N 32 --frag=no --splice -G 30000 -C 0 -u n --splice-flank=no -A 1 -B 2 -O 2,24 -E 1,0 -z 400,200 --no-end-flt. Reads were assigned to each canonical sgRNA by selecting those which overlapped the genomic interval 54-86 (leader), did not overlap the genomic interval 100-21546 (ORF1ab), and contained the specific downstream transcription regulatory sequence preceding each ORF (see Supplementary Table 2 for list of genomic intervals). Mapped reads filtering and counting were carried out by using Samtools version 1.9 (http://htslib.org).

#### Sequence alignments

The sequence alignments were realized using the Clustal Omega software (https://www.uniprot.org/align/).

### Statistical analysis

For the real-time PCR assays, Cq values of N1, N2 and N3 are reported as means ±standard deviation, as the ratios to the internal control detected with the CLONIT quanty COVID-19 kits (Ref. RT-25; IVD approved). Statistical significance was defined as a p value <0.05 *versus* the vehicle control. Immunofluorescence data were analysed by two-sample t-tests using the SPSS software, version 16.0. A probability (P)□<0.05 was considered statistically significant.

### Electrostatic potential and docking experiments

Electrostatic potential calculations were carried out with Adaptive Poisson-Boltzmann Solver software (41) within the AutoDock Tools environment (42) using the default parameters. Mapping of the electrostatic potential onto the protein surface and display of the molecules was carried out with PyMOL Molecular Graphics system (Version 1.8, Schrodinger LLC, 2015).

The PolyP molecule (ligand) containing 20 phosphate atoms (PolyP20) was built using the tools available within the ZINC data bank web server (43), and then translated into PDB coordinates. The three-dimensional structures of the receptor molecules were downloaded from the PDB. The atom types and charges were assigned to the ligands and receptors using the AutoDock Tools resources. All of the docking experiments were carried out with Autodock Vina (44).

## Supporting information

Supplemental Informations

## Data availability

*Gene sequences accession numbers:*

*GISAID: https://www.epicov.org/*
- 1-hCoV-19/Italy/CEINGE-Naples1/2020: EPI_ISL_477204 (entire virus sequence)
- 1-hCoV-19/Italy/CEINGE-Naples1/2020: EPI_ISL_514432 (entire virus sequence)
- hCoV-19/Italy/CEINGE-Naples2.1/2020: EPI_ISL_481510 (ORF1ab polyprotein, partial cds)
- hCoV-19/Italy/CEINGE-Naples2.2/2020: EPI_ISL_481511 (ORF1ab polyprotein, partial cds)
- hCoV-19/Italy/CEINGE-Naples3.1/2020: EPI_ISL_481512 (ORF1ab polyprotein, partial cds)
- hCoV-19/Italy/CEINGE-Naples2.3/2020: EPI_ISL_481716 (Gene S surface glycoprotein, partial cds)
- hCoV-19/Italy/CEINGE-Naples3.2/2020: EPI_ISL_481741 (ORF1ab polyprotein, partial cds)
-hCoV-19/Italy/CEINGE-Naples3.3/2020: EPI_ISL_481759 (Partial sequence of Spike glycoprotein gene S)
-hCoV-19/Italy/CEINGE-Naples4.1/2020: EPI_ISL_481760 (ORF1ab polyprotein, partial cds)
-hCoV-19/Italy/CEINGE-Naples4.2/2020: EPI_ISL_481761(ORF1ab polyprotein, partial cds)
-hCoV-19/Italy/CEINGE-Naples4.3/2020: EPI_ISL_481762 (Partial sequence of Spike glycoprotein gene S is reported).
- BetaCov/Korea/KCDC03/2020, registered ID: EPI_ISL_407193 (entire virus sequence)

Direct RNA sequencing using Nanopore Technology, data available at:

- http://www.ceinge.unina.it/polyp/ (Username: Covid19; Password: PolyP120)

## GenBank

MT682732.1 (Severe acute respiratory syndrome coronavirus 2 isolate SARS-CoV-2/human/ITA/Naples/2020, complete genome)

MT890669 (Severe acute respiratory syndrome coronavirus 2 isolate SARS-CoV-2/human/ITA/Naples/2020, complete genome)

## Acknowledgements

We would like to thank Prof. David Schlessinger Scientist Emeritus Intramural Research Program Ageing NIH Bethesda Maryland, USA for critical discussion of the manuscript, Dr. Alessandro Cometta ZEISS, Milan Italy for assistance on Zeiss Elyra 7 images. The authors thank further Dr. Antonio Limone (IZSM Director), Prof. Pietro Forestieri and Dr. Mariano Giustino (CEINGE President and CEO, respectively) for collaborative supporting the program within Regione Campania Covid19 Taskforce. The authors thank Prof.Ivan Gentile (Head of the Covid19 therapy Unit at AOU Federico II) for comments in clinical trials of Covid19 affected patients. This study was supported by the project “CEINGE TASK-FORCE COVID19”, code D64I200003800 by Regione Campania for the fight against Covid-19 (DGR n. 140 del 17 marzo 2020). The authors further thank the following resources agencies for grant support: the Italian Association for Cancer Research (AIRC) Grant IG 2219 (2018-2023; MZ), Fondazione Celeghin Italiana 2019-2020 (MZ), PRIN2017-2020-2023: Prot. 2017FNZRN3-LS2 (MZ). FISR-MIUR 03311-2020 (MZ). NRF 2018R1A5A2025079 - Korean Ministry of Science and ICT (J-HC).

## Author Contributions

V.F. performed RT-PCR analyses, F.A. performed quantitative immunofluorescence staining, R.S. prepared RNA from cell lysates and assemble the manuscript draft, L.M. performed immunoblotting, G.C. performed cell index assay. V.F., F.A. and L.M. performed in vitro treatment on not infected cells. V.F. and L.M. performed Caspase assays. F.A. amplified viral sequences for sequencing analyses. M.Z., G.F., M.V., S.B., G.B., P.C. and B.M.P. performed SARS-CoV-2 infections in BLS3 authorized lab. G.C. and M.C. obtained human primary cells from nasal brushing. G.P. and A.B. analyzed viral sequencing and bioinformatic analyses. L.C. and R.D.M. performed Nanopore sequencing. S.P. and M.B. performed docking experiments. C.T. and L.A. obtained samples from Italian COVI-19 patients. I.P. and B.I. performed staining on Vero cells. D.Y.K., K.S.Y., J.H.C, H.Y.K. performed experiments in Korean lab. M.Z. J.H.C, H.Y.K. designed the experiments, critical discuss with authors the experimental plans and wrote the paper. All authors discussed the results and commented on the paper. All other authors declare no competing interests.

---

**Supplementary Fig. 1.**
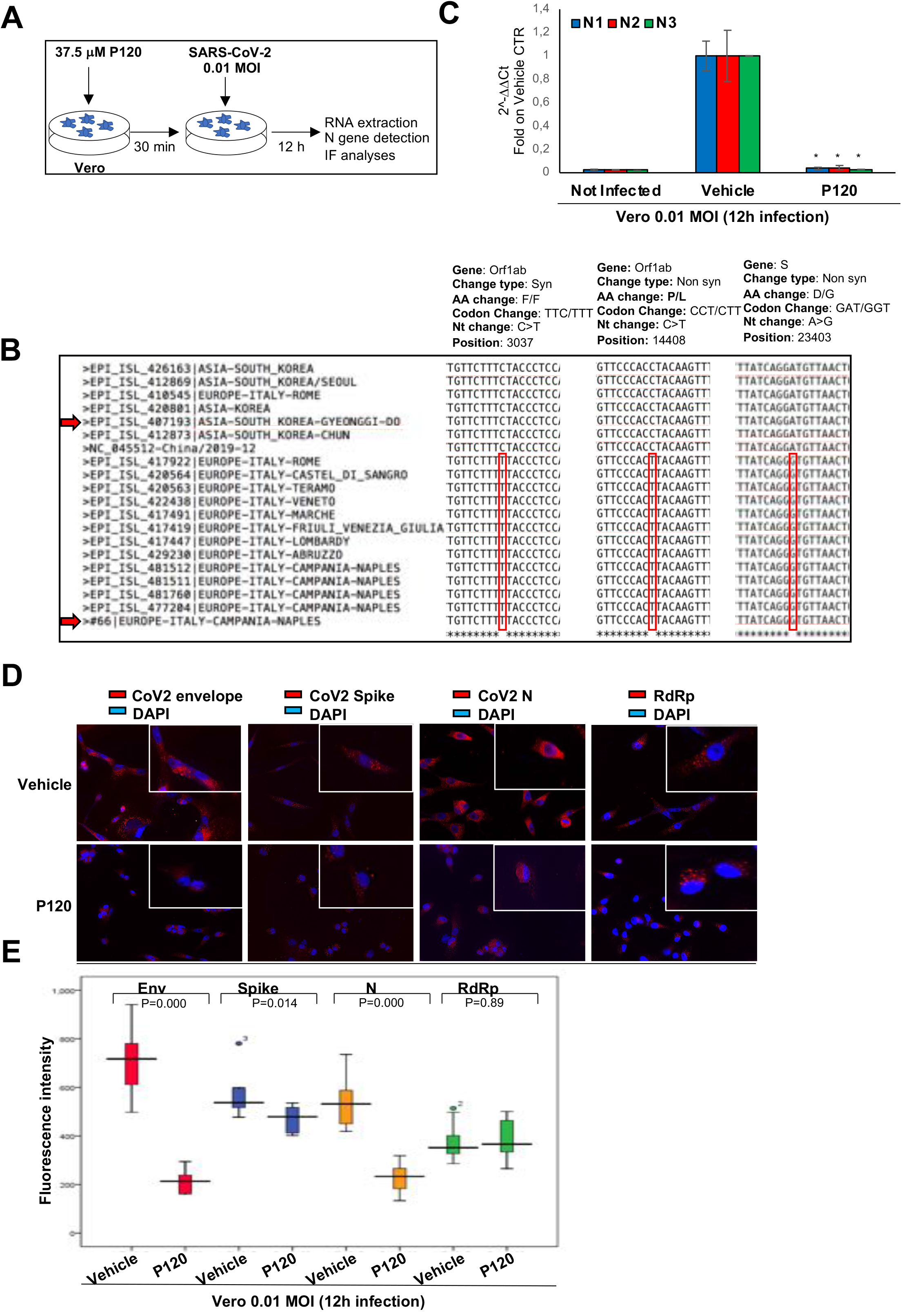

**Supplementary Fig. 2.**
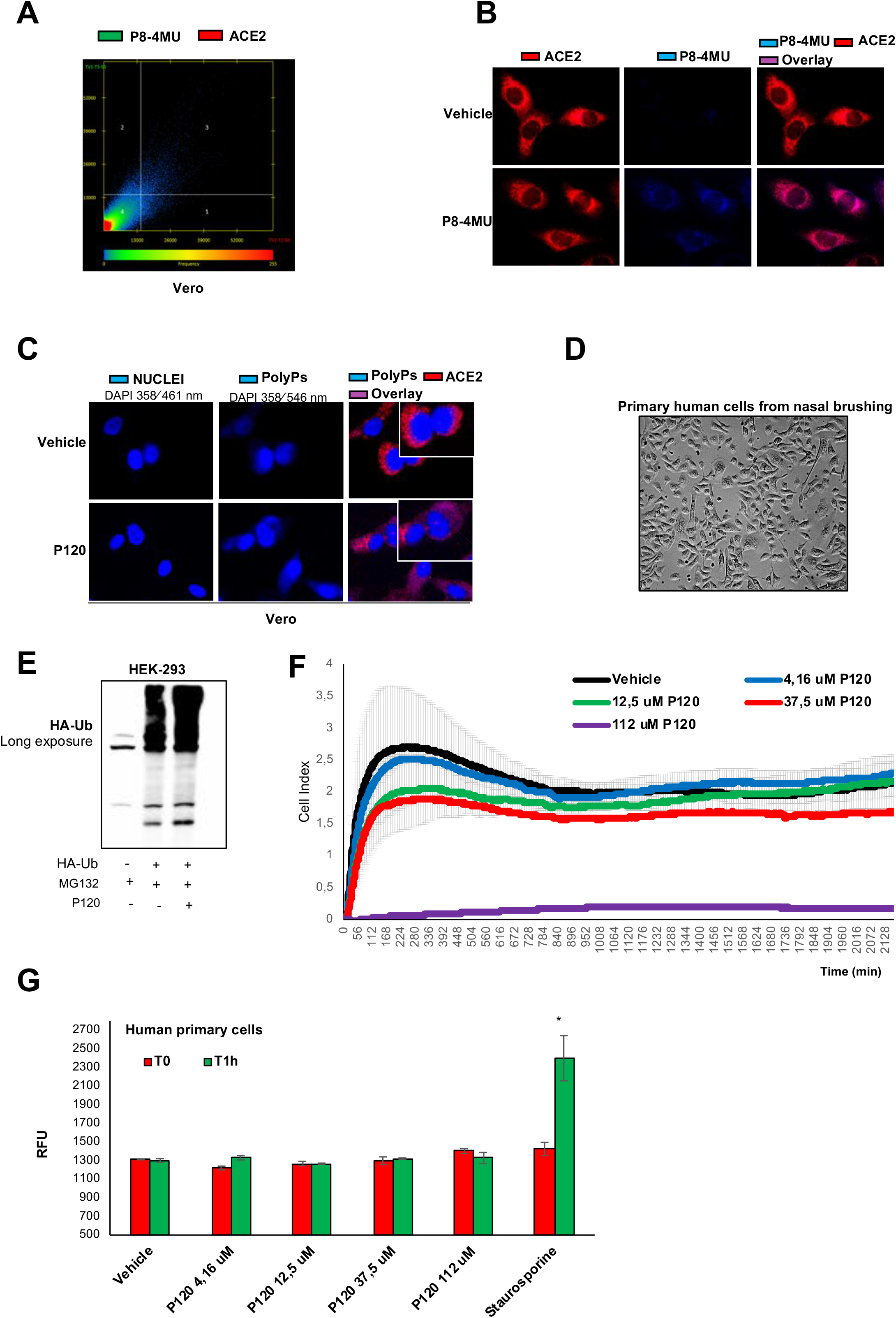

**Supplementary Fig. 3.**
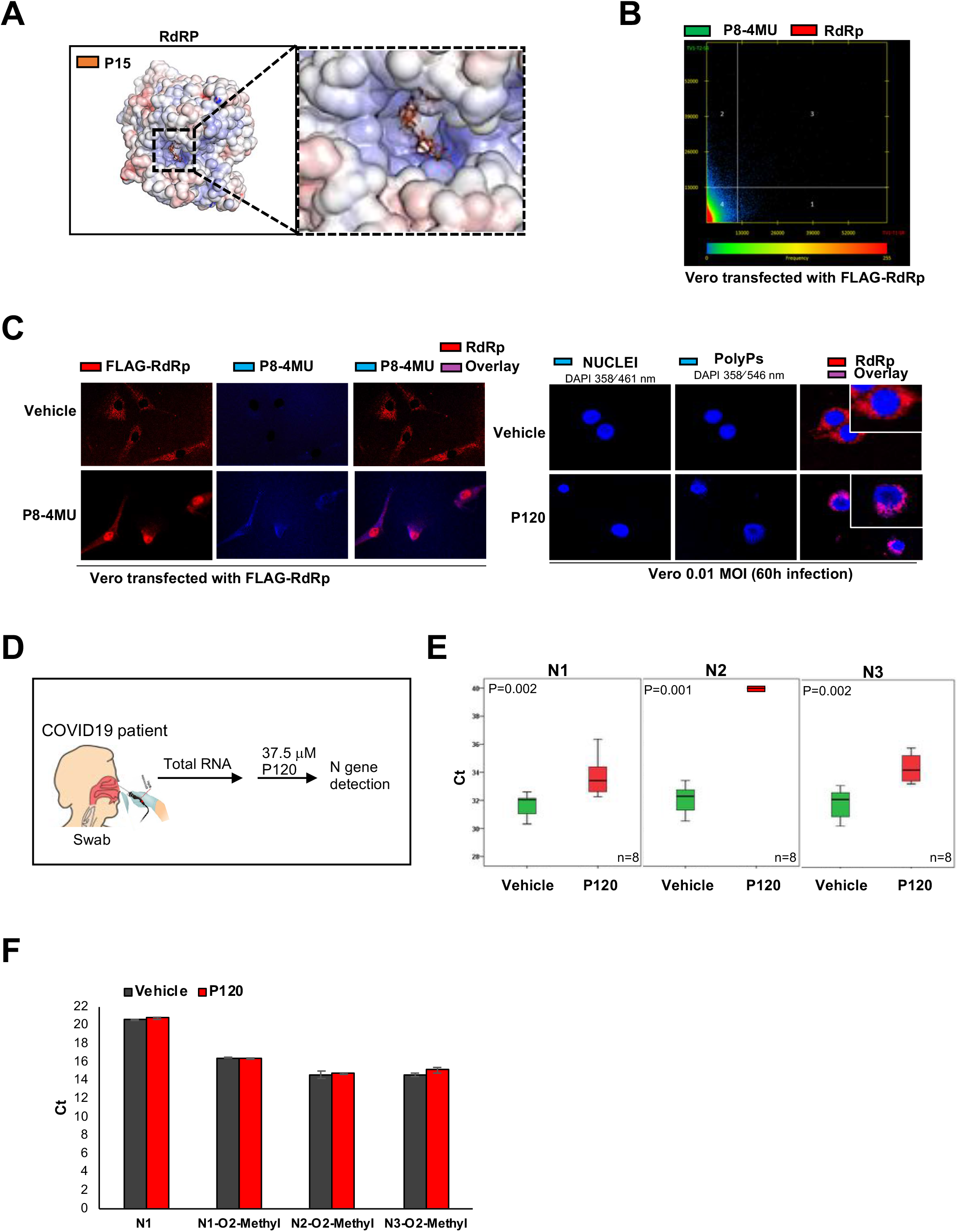

**Supplementary Fig. 4.**
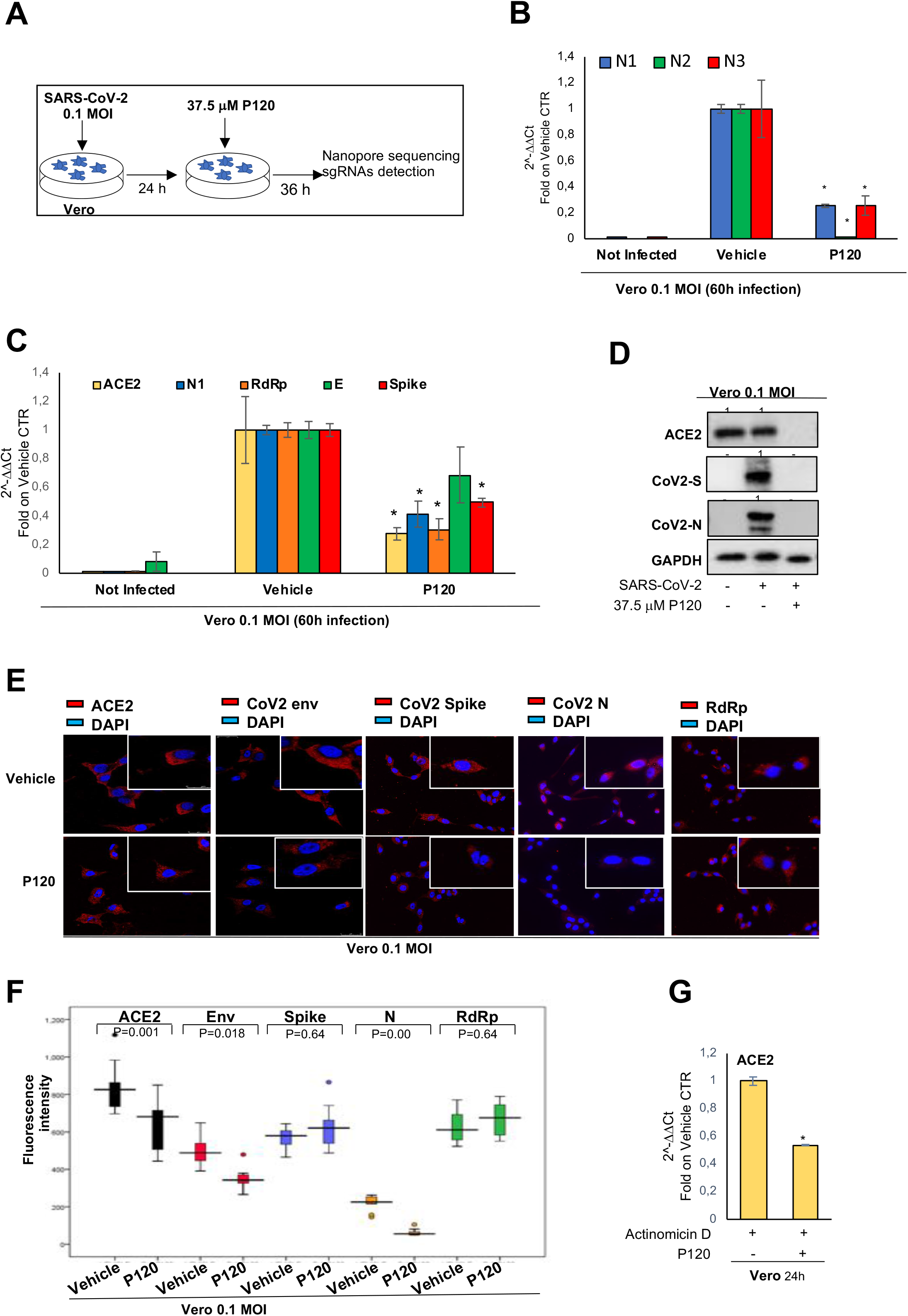

**Supplementary Fig. 5.**
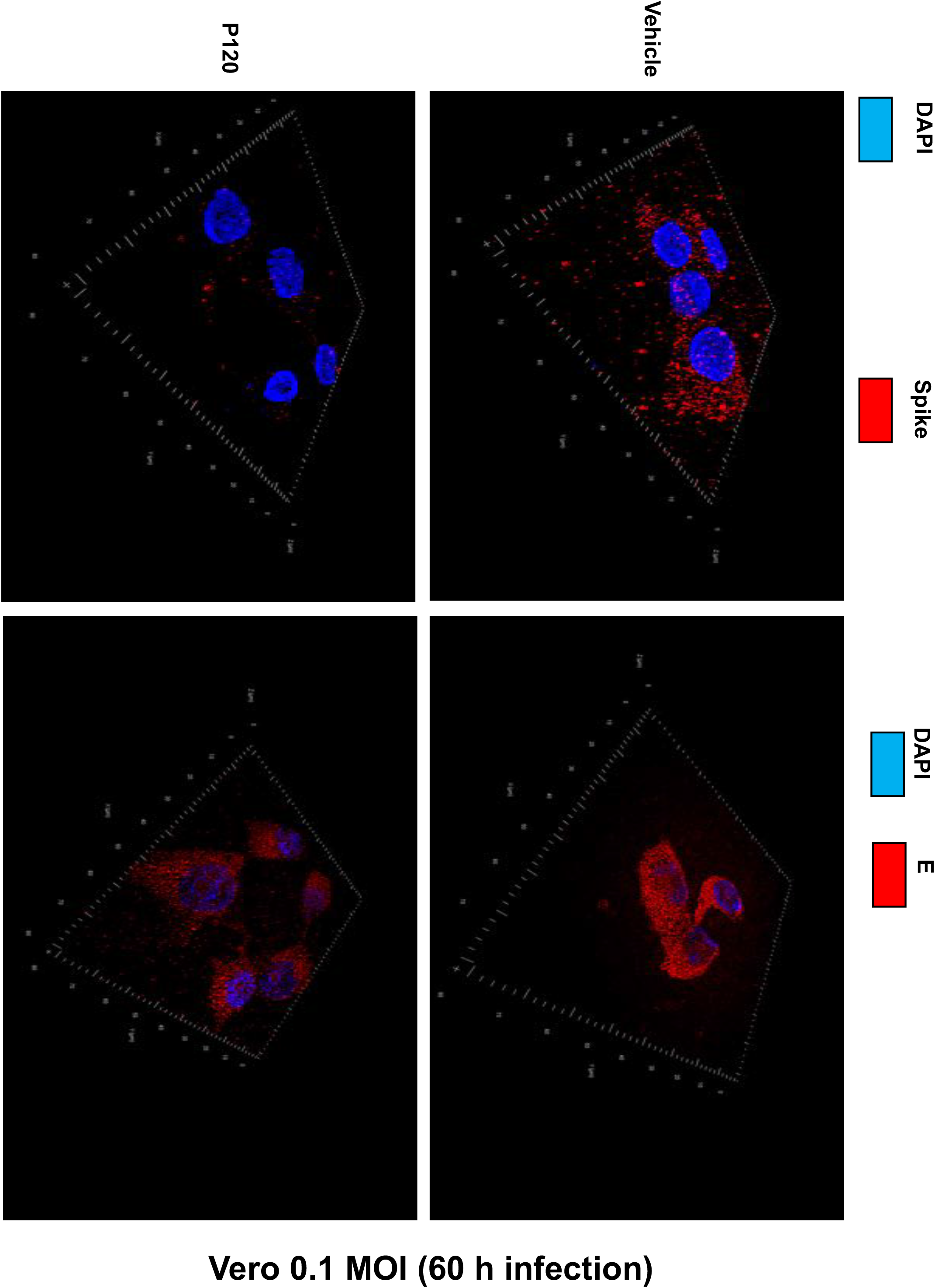

**Supplementary Fig. 6.**
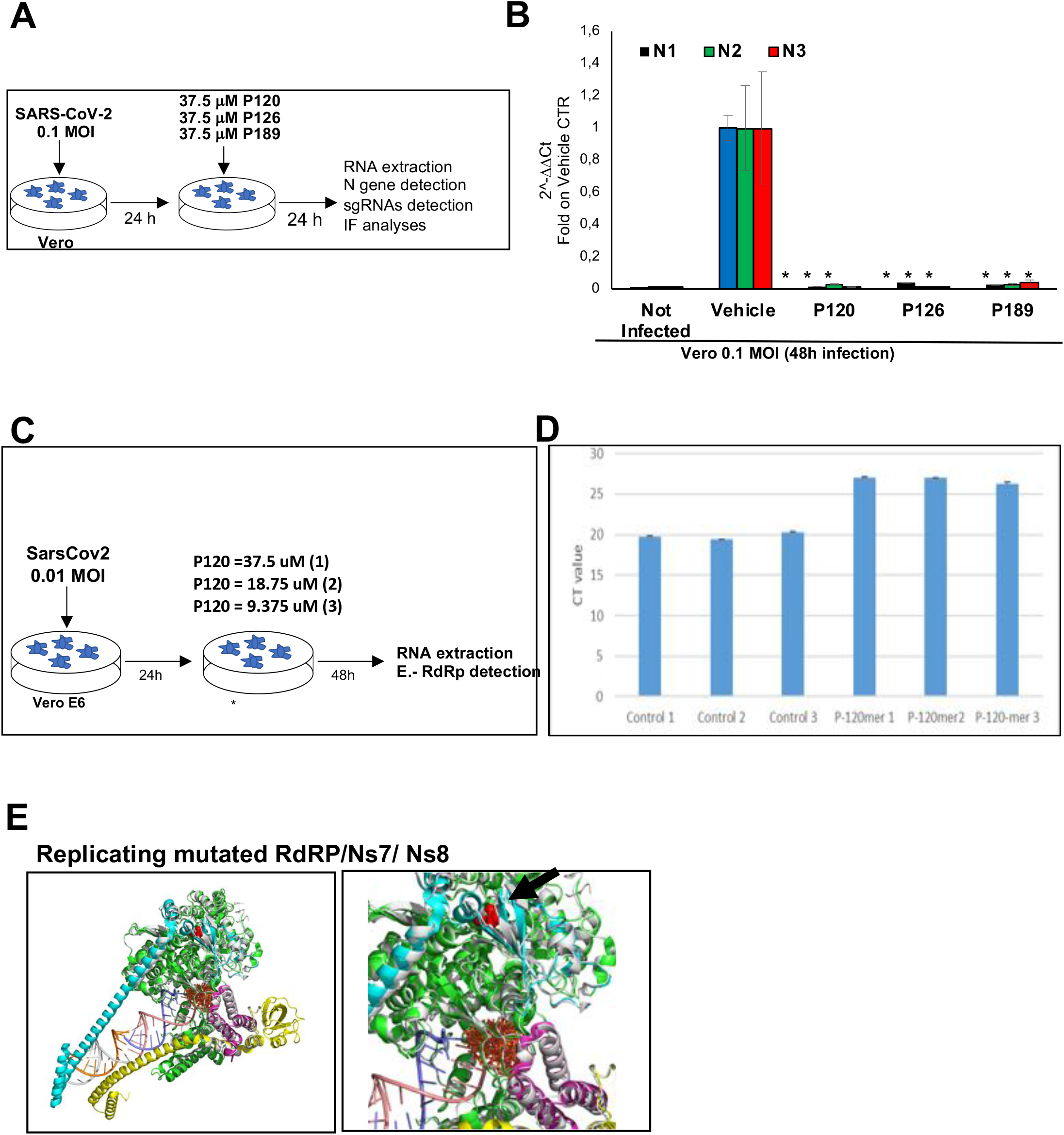

**Supplementary Fig. 7.**
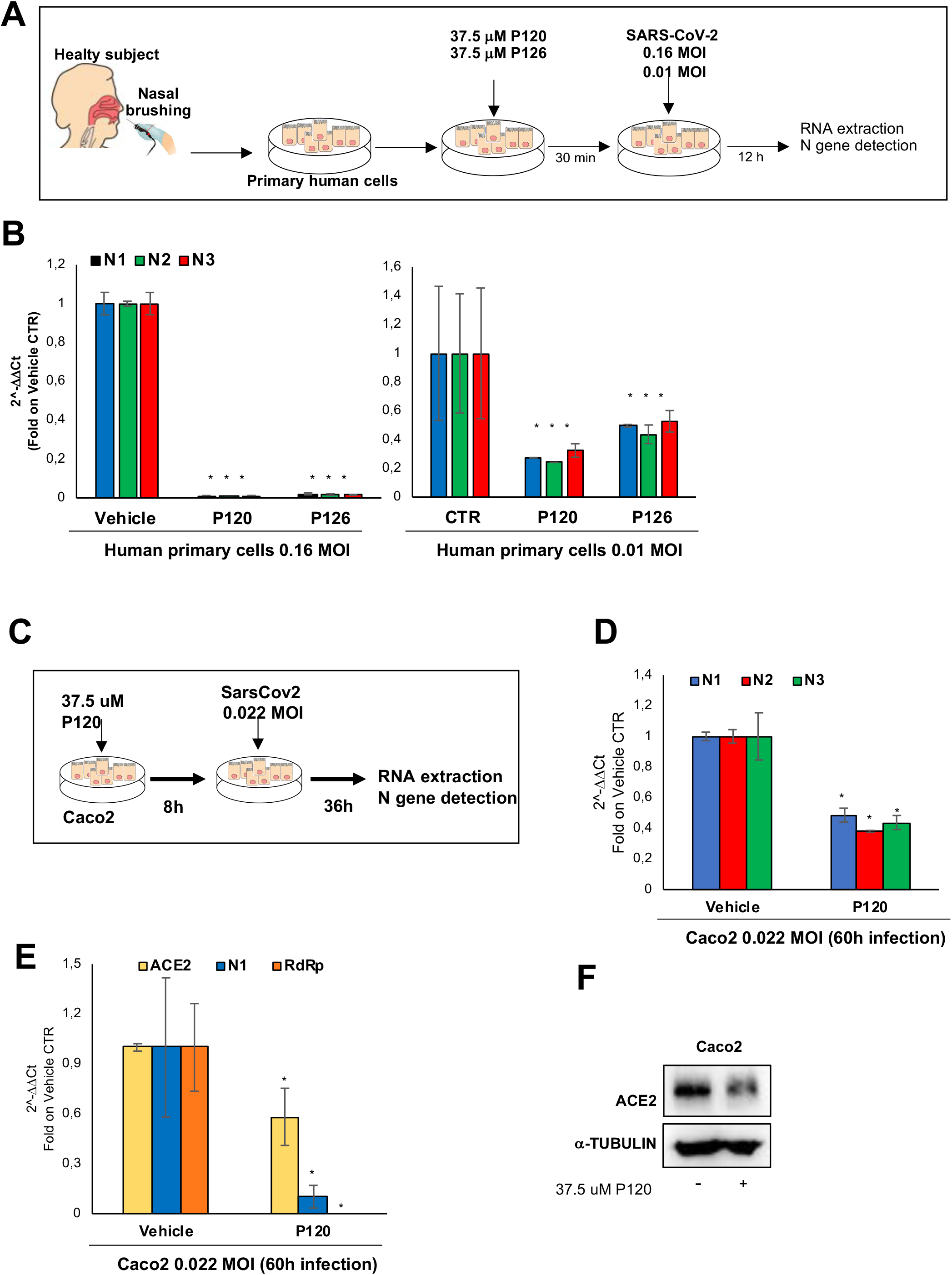

**Supplementary Fig. 8.**
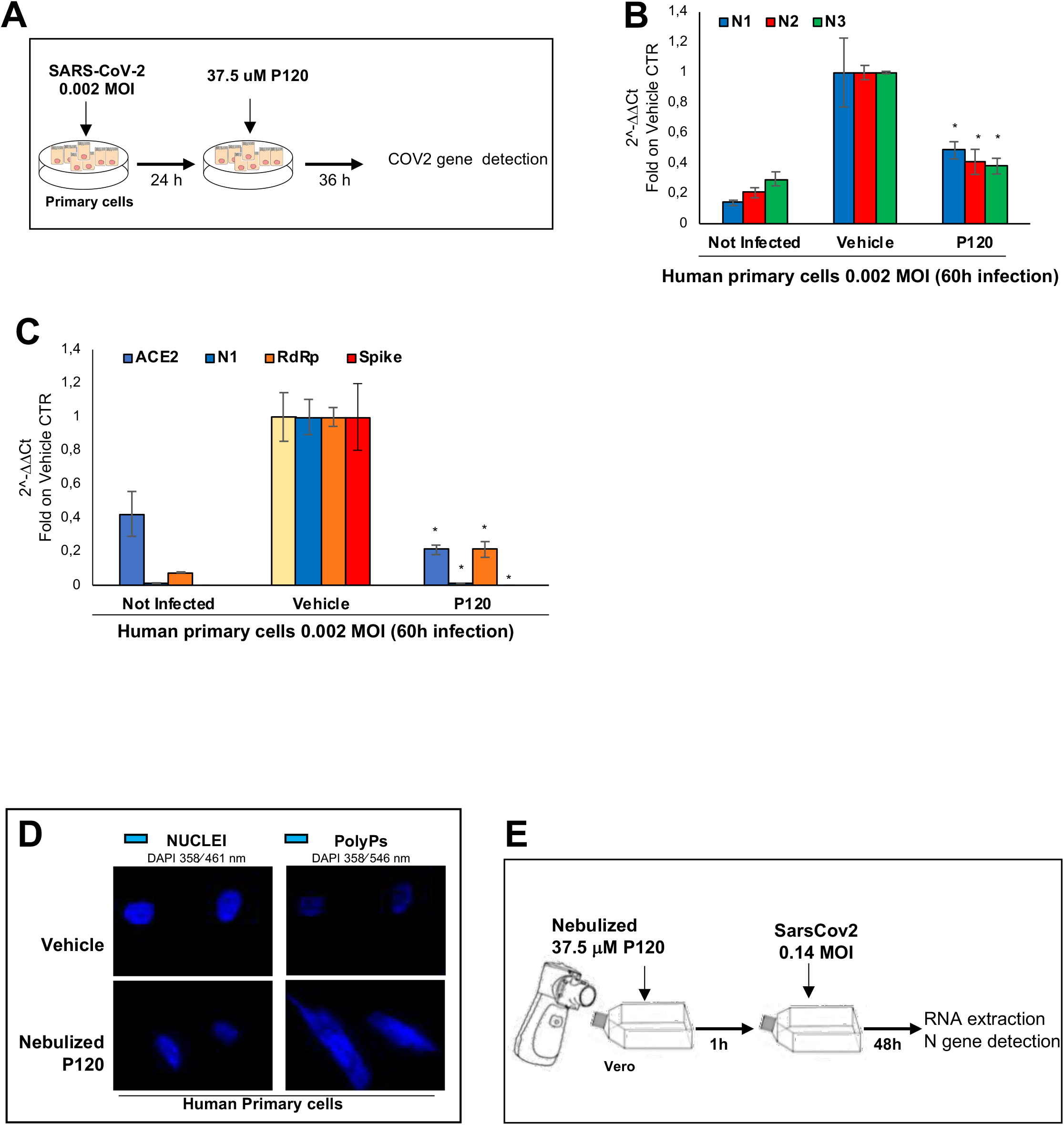

## Notes

### Competing Interest Statement

The authors have declared no competing interest.

