## Supplemental Informations for "Long-chain polyphosphates impair SARS-CoV-2 infection and replication: a route for therapy in man"

5 U.O.C. di Patologia Clinica Ospedale D. Cotugno, Azienda Sanitaria Ospedali dei Colli, Naples, Italy.

6 Università La Sapienza di Roma, Rome, Italy

7 HAIM BIO Co. Ltd, Industrial Park, Korea University, 145 Anam-ro, Seongbuk-gu, Seoul, South Korea

8 Department of Surgery, Yonsei University College of Medicine, Seoul, Korea

***Corresponding authors:**

Jae-Ho Cheong –

Hong-Yeoul Kim –

Massimo Zollo –

**The PDF file includes:**

**Fig. S1 to S8**

**Tables S1 and s2**

**Supplementary Materials and Methods**

**Fig. S1. Anti-viral effects against SARS-CoV-2 of ‘preventive treatments’ with long-chain PolyPs. (A)** Experimental plan. Vero cells were treated with 37.5 μM PolyP120. After 30 min, the cells were infected with SARS-CoV-2 viral particles (0.01 MOI). After 12 h, cells were lysed and RNA and protein were extracted. Non-infected cells were used as negative control. **(B)** Alignment analyses of the SARS-CoV-2 sequences from South Korea and Italy, showing synonymous and non-synonymous variants for ORF1ab (i.e., RdRp gene) and non-synonymous variants for S genes in the Italian sequences (which can affect proofreading activity of RdRp protein(1)). **(C)** Ct values from real-time RT-PCR following RNA extraction, showing quantification of expression of N1, N2 and N3. **(D-E)** Immunofluorescence was with antibodies against the S, E, N and RdRp proteins. The image acquisition and the intensity of staining was analyzed with MANTRA quantitative workstation pathology. The fluorescence intensity was measured in each cell and compared to vehicle control, with more than 100 cells counted. Representative images **(D)** and quantification of expression levels **(E)** are shown. Magnification 40×.

**Fig. S2. Anti-viral effects of PolyP120 are mediated by ACE2 down-regulation, with no cytotoxic effects. (A)** Immunofluorescence was performed on Vero cells treated with 37.5 μM PolyP8 conjugated with 4-methylumbelliferone (P8-4MU) and the vehicle control, by using anti-ACE2 antibody. Image acquisition was made with ZEISS Elyra 7 with the optical Lattice SIM technology by using 63x oil immersion objective. The results of fluorescence colocalization between PolyPs (i.e., P8) and ACE2 are graphically represented as scatterplots where the distribution of the intensity of one color (i.e., green for P8) is plotted against that of the second color (i.e., red for ACE2) for each pixel. The number of the pixels is shown as “frequency” (0-255). Colocalization scatterplot is obtained by using ZEISS ZEN software (blue edition). **(B)** Vero cells treated with 37.5 μM PolyP8 conjugated with 4-methylumbelliferone (P8-4MU), and the vehicle control, were fixed and immunofluorescence was performed using an anti-ACE2 antibody. Acquisition was made with MANTRA Quantitative Pathology workstation. Magnification 40×. **(C)** SARS-CoV-2–infected Vero cells treated with 37.5 μM P120, and the vehicle control, were fixed and immunofluorescence was performed using an anti-ACE2 antibody. PolyPs were detected using DAPI (laser, 358/546 nm). Acquisition was made with MANTRA Quantitative Pathology workstation. Magnification 40×. **(D)** Human primary nasal epithelial cells obtained from nasal brushing of three healthy subjects. Magnification 40×. **(E)** Plasmid containing HA-tagged Ubiquitin was transiently expressed in HEK-293 cells. After 24h from transfection cells were treated with 10 μM MG132 (proteasome inhibitor) and 37.5 μM PolyP120 for additional 24h. Vehicle-treated cells with MG132 and non-trasfected cells were used as negative controls. Immunoblotting was performed with antibody against HA-Ubiquitin. Long exposure is shown. **(F)** Real-time cell proliferation analysis for the Cell Index approach. Human primary nasal epithelial cells (8,000) were plated and treated with PolyP120 (as indicated), with vehicle-treated cells as negative control. Impedance was measured every 2 min over 36 h (2160 min). Cytotoxicity was seen for 112 μM PolyP120. **(G)** Apoptosis assay using caspase-3 activity assay kits. Human primary nasal epithelial cells (300,000) were plated and treated with PolyP120 (as indicated), with vehicle-treated cells as negative control, and staurosporine (1 μM) as positive control. Fluorescence were measured before (T0) and after 1 h (T1h). No apoptosis was seen for PolyP120-treated cells.

**Fig. S3. PolyP120 inhibits transcription of viral sgRNAs through inhibition of RdRp. (A)** Molecular docking of PolyP15 on SARS-CoV-2 RdRp (corresponding to the PDB structure 6M71). RdRp is represented as a molecular surface coloured according to electrostatic potential. Colour scale ranges from -10 kT/e (red) to +10kT/e (blue). The ligand 15P is represented as orange sticks (left). Expanded view (right). **(B)** Immunofluorescence was performed on Vero cells transfected with FLAG-RdRp and treated with 37.5 μM PolyP8 conjugated with 4-methylumbelliferone (P8-4MU), and the vehicle control, by using an anti-RdRp antibody. Image acquisition was made with ZEISS Elyra 7 with the optical Lattice SIM technology by using 63x oil immersion objective. The results of fluorescence colocalization between PolyPs (i.e.,P8) and RdRP are graphically represented as scatterplots where the distribution of the intensity of one color (i.e., green for P8) is plotted against that of the second color (i.e., red for RdRP) for each pixel. The number of the pixels is shown as “frequency” (0-255). Colocalization scatterplot is obtained by using ZEISS ZEN software (blue edition). **(C)** **(Left panel)** Vero cells transfected with FLAG-RdRp and treated with 37.5 μM PolyP8 conjugated with 4-methylumbelliferone (P8-4MU), and the vehicle control, were fixed and immunofluorescence was performed using anti-RdRp antibody. Images are acquired using MANTRA quantitative workstation pathology. Magnification 40x. **(Right panel**) Vero cells were infected with SARS-CoV-2 and treated with 37.5 μM PolyP120 or vehicle control, and fixed. Immunofluorescence was performed with an antibody against RdRp, and PolyP120 detection with DAPI. Magnification 63×. (**D)** Experimental plan. For *in-vitro* experiments, RNA samples were obtained from frozen swabs from patients positive for COVID-19 (0.09 MOI) and treated with 37.5 μM PolyP120 or vehicle control. **(E-F)** Quantitative real-time reverse transcription assays for effects of PolyP120 on N1, N2 and N3 fragments from the N gene (from 8 patients) **(E)**, and amplification of N1 when external Spike RNA controls (N1/2/3-O_2_-methyl) were used **(F)**.

**Fig. S4. Therapeutic treatments with long-chain PolyPs*.* (A)** Experimental plan. Vero cells (425,000) were infected with SARS-CoV-2 viral particles, with non-infected cells as negative control of infection. After 24 h, cells were treated with 37.5 μM P120, and 36 h later they were lysed and RNA was extracted. **(B)** Ct values from real-time RT-PCR following RNA extraction, showing quantification of expression of N1, N2 and N3 (0.1 MOI). **(C)** Ct values from real-time RT-PCR are shown for expression of the ACE2, N1, E, RdRp, S genes, as indicated (0.1 MOI). **(D)** Immunoblotting of non-infected and SARS-CoV-2–infected Vero cells treated with 37.5 μM PolyP120 and vehicle, using antibodies as indicated (0.1 MOI). **(E-F)** Immunofluorescence with antibodies against the S, E, N and RdRp proteins. Images acquisition and intensity of staining were analyzed with MANTRA Quantitative workstation pathology. Fluorescence Intensity was measured in each cell and compared to vehicle control, with more than 100 cells counted. Representative images **(E)** and quantification of expression levels **(F)** are shown. Magnification 40×. **(G)** Vero cells (500,000) were plated and treated for 24 h with 10 μg/mL actinomycin D (transcription inhibitor) or vehicle, and 37.5 μM PolyP120. Ct values from real-time RT-PCR following RNA extraction, showing quantification of expression of ACE2.

**Fig. S5**. **Therapeutic treatments with PolyP120 in cells*.*** Immunofluorescence with antibodies against S and E proteins on Vero cells infected with SARS-CoV-2 viral particles and treated with PolyP120. The SIM image was acquired with Elyra 7 and 3-dimentional (3D) reconstruction of Z stacking data were performed by using ZEISS ZEN software (blue edition). Magnification 63×.

**Fig S6. Therapeutic efficacy of PolyP120 in different SARS-CoV-2 viral strain*.* (A)** Experimental plan. Vero cells (425,000) were infected with SARS-CoV-2 viral particles (0.09 MOI), with non-infected cells as negative control of infection. After 24 h, cells were treated with 37.5 μM PolyP120, PolyP126 or PolyP189, and 24 h later they were lysed and RNA was extracted. **(B)** Ct values from real-time RT-PCR following RNA extraction, showing quantification of expression of N1, N2 and N3 following PolyPs treatments. **(C)** Experimental plan. Vero cells (425,000) were infected with SARS-CoV-2 viral particles (0.01 MOI) from a Korean patient positive for SARS-CoV-2, with non-infected cells as negative control of infection. After 24 h, cells were treated with 9.38, 18.75 and 37.5 μM PolyP120, and 24/48 h later they were lysed and RNA was extracted. **(D)** Ct values from real-time RT-PCR following RNA extraction, showing quantification of expression of E and RdRp following the treatments with the different PolyP120 concentrations. **(E)** Molecular docking between mutated RdRp/nsp7/nsp8 and PolyP15. RdRp mutation at position 14,408 is shown in red (black arrow). Expanded view (right).

**Fig. S7. Preventive efficacy of PolyP120 in SARS-CoV-2 infected human cells. (A)** Experimental plan. Primary nasal epithelial cells (600,000) were treated with 37.5 μM P120 or P126. After 20 min, these pretreated cells were infected with SARS-CoV-2 viral particles (0.16-0.01 MOI), with non-infected cells as negative control of infection. After 12 h, cells were lysed and RNA was extracted. **(B)** Ct values from real-time RT-PCR following RNA extraction, showing quantification of expression of N1, N2 and N3 following treatments with PolyP120 and PolyP126 at 0.16 (left) and 0.01 (right) MOI. **(C)** Experimental plan. Caco2 cells (500,000) were treated with 37.5 μM PolyP120. After 8 h, these pretreated cells were infected with SARS-CoV-2 viral particles (0.022 MOI), with non-infected cells as negative control of infection. After 36 h, these Caco2 cells were lysed and RNA was extracted. **(D-E)** Ct values from real-time RT-PCR following RNA extraction, showing quantification of expression following treatment with PolyP120 of N1, N2 and N3 **(D)** and ACE2, N1 and RdRp **(E)**. **(F)** Immunoblotting on Caco2 cells treated with 37.5 μM PolyP120 and vehicle, using antibodies as indicated.

**Fig. S8. Therapeutic efficacy of PolyP120 in SARS-CoV-2 infected human cells. (A)** Experimental plan. Human primary nasal epithelial cells (425,000) were infected with SARS-CoV-2 viral particles (0.002 MOI), with non-infected cells as negative control of infection. After 24 h, cells were treated with 37.5 μM PolyP120, and 36 h later they were lysed and RNA was extracted. **(B-C)** Ct values from real-time RT-PCR following RNA extraction, showing quantification of expression of N1, N2 and N3 **(B)** and ACE2, N1, RdRp and S **(C)** following treatments with PolyP120. **(D)** Vero cells treated with nebulized 37.5 μM P120, and the vehicle control, were fixed and immunofluorescence was performed. PolyPs were detected using DAPI (laser, 358/546 nm). Acquisition was made with MANTRA Quantitative Pathology workstation. Magnification 40×. **(E)** Experimental plan. Vero cells (400,000) were treated with nebulised 37.5 μM PolyP120. After 1 h, these cells were infected with SARS-CoV-2 viral particles (0.14 MOI), with non-infected cells as negative control of infection. After 48 h, the cells were lysed and RNA was extracted.

**Table S1.** Oligonucleotide sequences of the primers used in this study.

| **Target viral gene** | **Primer** | **Primer sequence** |
| --- | --- | --- |
| ACE2 | Forward | GAAATTCCCAAAGACCAGTGGA |
|  | Reverse | CCCCAACTATCTCTCTCGCTTCAT |
| RdRp | Forward | GTGAAATGGTCATGTGTGGCGG |
|  | Reverse | CAAATGTTAAAAACACTATTAGCATA |
| N1 | Forward | GACCCCAAAATCAGCGAAAT |
|  | Reverse | TCTGGTTACTGCCAGTTGAATCTG |
| Spike | Forward | ATTGCCACTAGTCTCTAGT |
|  | Reverse | AGGATCTGAAAACTTTGTCA |
| Envelope | Forward | ACAGGTACGTTAATAGTTAATAGCGT |
|  | Reverse | ATATTGCAGCAGTACGCACACA |
| sg (2) | Forward | CAAACCAACCAACTTTCGATCTCTTGTA |
| sgS | Reverse | TGAAAGAATTAGTGTATGCA |
| sgE | Reverse | AGAAGTACGCTATTAACTATT |
| sgM | Reverse | TATTACTAGGTTCCATTGTTCAA |
| sgN | Reverse | TCTGGTTACTGCCAGTTGAATC |
| Interleukin-6 | Forward | GCCACTCACCTCTTCAGAAC |
|  | Reverse | AGCATCCATCTTTTTCAGCC |
| Interleukin-10 | Forward | CCTGCCTAACATGCTTCGAGA |
|  | Reverse | TGTCCAGCTGATCCTTCATTTG |
| Interleukin-12 | Forward | TGATGGCCCTGTGCCTTAGT |
|  | Reverse | GGATCCATCAGAAGCTTTGCA |
| Interferon-γ | Forward | AGGCATTTTGAAGAATTGGAAAGA |
|  | Reverse | AGTAAAAGGAGACAATTTGGCTCT |
| Tumour necrosis factor-α | Forward | TCTCTCTAATCAGCCCTCTGG |
|  | Reverse | GCTACATGGGCTACAGGC |

**Table S2.** Reads assigned to sgRNA (per 1000 reads assigned to host genome).

| **sgRNA** | **Vehicle** | **PolyP120** |
| --- | --- | --- |
| sG-S | 0.49 | 0.04 |
| sG-ORF3a | 7.05 | 1.77 |
| sG-E | 4.01 | 1.22 |
| sG-M | 38.67 | 7.09 |
| sG-ORF6 | 14.00 | 1.85 |
| sG-ORF7a | 71.18 | 7.27 |
| sG-ORF7b | 0.29 | 0.00 |
| sG-ORF7b-short | 2.06 | 0.30 |
| sG-ORF8 | 34.66 | 3.25 |
| sG-N | 592.42 | 40.60 |
| sG-ORF10-short | 0.78 | 0.00 |

**Supplementary material and methods**

***Preparation of sodium polyphosphate***

*Synthesis of sodium polyphosphate glass*

Sodium phosphate monobasic (NaH_2_PO_4_ ≥99.0%; Sigma Aldrich) was polymerised in an aluminium crucible in an electric furnace (HQ-DMF3; Coretech, Korea) at 700 ℃ for 1 h. The molten sodium polyphosphate was then poured onto a copper plate for rapid cooling, and the sodium polyphosphate glass obtained was ground into a powder.

*Fractionation of the polyphosphates*

In general, the synthesised polyphosphate polymer has a wide range of chain lengths. Fractionation of the polyphosphates (PolyPs) was carried out by fractional precipitation. The sodium polyphosphate powder (20 g) was dissolved in 200 mL distilled water (i.e., 1:10, w/v), the pH was adjusted to 7, and it was left at 25 ℃ for 12 h. Acetone was then added to this solution of sodium polyphosphate drop-wise until the solution became cloudy due to PolyPs precipitation. The precipitated PolyPs were separated by centrifugation at 4200 rpm for 5 min (1248R; Labogene, Denmark). This fractional precipitation was repeated 25 times, with the PolyPs fractions obtained dried in a freeze drier for 48 h.

*Determination of PolyPs chain length*

A gel permeation chromatography–multi-angle light scattering system was used to determine the PolyPs chain lengths in each of the precipitation fractions. This included an HPLC system (LC-20AD; Shimadzu, Japan) and a multi-angle light scattering detector (miniDawn Treos II; Wyatt Technology, USA). The gel permeation chromatography column (PL aquagel-OH; 7.5 × 50 mm; 8 µm; Agilent Technology, USA) was protected by a guard column (PL aquagel-OH Mixed-M; 7.5 × 300 mm; 8 µm; Agilent Technology, USA). The system was run with 150 mM NaCl. The concentration of PolpPs in the sample injection was 25 mg/mL, and the injection volume was 50 µL. The temperature of the column was maintained at 30 ℃.

*Production of Polyphosphates*

Polymerisation of sodium phosphate monobasic takes place as follows:

*n*NaH_2_PO_4_ = (NaPO_3_)*_n_* + *n*H_2_O

The polymer obtained is known as Graham’s salt, which contains the polyphosphates (PolyPs) in three mixed molecular structures: linear, cyclic and branched. The branched PolyPs are very labile, and when they dissolve in water, the branching points of these PolyPs are hydrolysed at a high rate irrespective of pH, and even at room temperature. In contrast, the linear PolyPs and cycloPolyPs are hydrolysed very slowly at neutral pH and room temperature. The ‘half hydrolysis time’ for the P–O–P bonds in linear PolyPs at pH 7 at 25 °C is several years. These synthesised polyphosphates usually contain very small amounts of cycloPolyPs, and almost all of these have chain lengths of <6 inorganic phosphate units (3) . In the present experiments, the minimum chain length obtained for these PolyPs was 20, and therefore, when these PolyPs were separated by fractional precipitation the cycloPolyPs were not precipitated.

After fractionation of the PolyPs by gel permeation chromatography, their chain lengths (n) were determined according to the following formula:

Mn (number-average molar mass) = ${Na}_{n+2}P_{n}O_{3n+1}$

The range of the chain lengths in each fraction were then obtained through the polydispersity index (weight-average molar mass/ number-average molar mass; Mw/Mn), using the Astra software, version 6.1 (Wyatt Technology, USA).

***Immunoblotting***

Cells for immunoblotting were washed in cold phosphate-buffered saline (PBS) and lysed in cell lysis buffer (20 mM sodium phosphate, pH 7.4, 150 mM NaCl, 10% [v/v] glycerol, 1% [w/v] sodium deoxycholate, 1% [v/v] Triton X-100) supplemented with protease inhibitors (Roche, Basel, Switzerland). The resulting cell lysates were cleared by centrifugation at 16,200× *g* for 10 min at room temperature, and the supernatants were removed and assayed for protein concentration using the Protein Assay Dye Reagent (BioRad). The cell lysate proteins (50 μg) were separated using SDS-PAGE gels of different percentages, which depended on the molecular weights of the proteins of interest. The proteins were then electrophoretically transferred to PVDF membranes (Millipore). After 1 h in blocking solution with 5% (w/v) dry milk fat in PBS, or 5% (w/v) bovine serum albumin (Sigma-Aldrich) in Tris-buffered saline (both of which contained 0.02% [v/v] Tween-20), the PVDF membranes were incubated with the required primary antibody overnight at 4 °C: anti-ACE2 (1:1000; ab15348), anti-SARS-CoV-2 Nucleoprotein (1:250; 35-579; ProSci Inc.), anti-SARS-COV2 Spike (1: 250; ab272504), anti-p-NFkB p65 (Ser 311) (1:500; sc-101748), anti-HA (1:500; 12CA5; Merck), anti-GAPDH (1:10 000; sc-365062), or anti-α-tubulin (1:3000; Ab15246), anti-β-actin (1:10000; A5441; Sigma). The membranes were then incubated with the required secondary antibodies for 1 h at room temperature: secondary mouse or rabbit horseradish-peroxidase-conjugated antibodies (NC 15 27606; ImmunoReagents, Inc.), diluted in 5% (w/v) bovine serum albumin in TBS-Tween or in 5% (w/v) milk fat in PBS-Tween, according to the manufacturer instructions. The protein bands were visualised using chemiluminescence detection (Pierce-Thermo Fisher Scientific Inc., IL, USA). Western blotting was performed in triplicate. Densitometry analysis was carried out using the ImageJ software. The peak areas of the bands were measured on the densitometry plots, and the percentages were calculated. Then, the density areas of the peaks were normalised with those of the loading controls, and the ratios for the corresponding controls are presented as fold-changes.

***Immunofluorescence and HuluFISH***

*Infected cells*

Vero cells (1 ×10^4^) were plated onto Collagen-1–coated glass coverslips in 24-well plates, and infected with SarsCov2 viral particles and treated with 37.5 μM PolyPs as described above.

*Non-infected cells*

Vero cells (1 ×10^4^) were plated onto Collagen-1–coated glass coverslips in 24-well plates, and treated with 37.5 μM P8-4MU for 12 h.

*Nebulised PolyP120 treatment*

Human primary nasal epithelial cells (1 ×10^4^) were plated onto Collagen-1–coated glass coverslips in 24-well plates, and treated with the nebulised 37.5 μM PolyP120 (dissolved in 2 mL; condensation rate: 1 mL/min) for 24 h.

*Cell processing*

The cells were fixed in 4% paraformaldehyde in PBS for 30 min. washed three times with PBS, and permeabilised with 0.1% Triton X-100 (215680010; Acros Organics) diluted in PBS, for 15 min. The cells were then washed with PBS and blocked with 3% bovine serum albumin (A9418; Sigma) in PBS for 1 h at room temperature. The samples were incubated with the relevant primary antibodies overnight at 4 °C: anti-ACE2 (1:50; ab15348; Abcam), anti-SARS Spike (1:50; ab272420; Abcam), anti-SARS-CoV RdRp R2 (1:50; ARG1109; Arigo Biolaboratories), anti-SARS-CoV-2 Nucleoprotein (1:50; 35-579; ProSci Inc.), or anti-SARS-Cov Envelope (1:50; IT-002-005; ProSci). After washing twice with PBS, the samples were incubated with the relevant secondary antibody at room temperature for 1 h: anti-mouse Alexa Fluor 647 (1:100; ab150115; Abcam) and anti-Rabbit Alexa Fluor 647 (1:100; ab150075; Abcam). DNA and PolyPs were stained with DAPI (#62254; Thermo Fisher). The slides were washed, and mounted with cover slips using 50% glycerol (G5150; Sigma-Aldrich).

HuluFISH was performed on Pan-SARS-CoV-2 probe against gRNA-Spike (MetaSystem – ZEISS) on SARS-CoV-2-infected Vero cells previously fixed in 4% paraformaldehyde in PBS and stained with antibody against ACE2 protein.

Microscopy image was carried out with ZEISS Elyra 7 with the optical Lattice SIM technology by using 63x oil immersion objective. Z stacking imaging, 3-dimentional (3D) reconstruction and colocalization scatterplots were performed by using ZEISS ZEN software (blue edition). The quantification of the fluorescence intensity was obtained with Mantra Quantitive Pathology Workstation, with the cells counted under immunofluorescence staining using the ImageJ software (version 1.52r; <https://imagej.nih.gov/ij/index.html>).

***Real-time cell proliferation assay***

The cell proliferation rates were determined using the xCELLigence system (Roche). Human primary nasal epithelial cells (5000 cells) were treated with increasing concentrations of PolyP120 and then harvested and washed with PBS, and resuspended in Dulbecco’s modified Eagle’s medium (DMEM) with 10% foetal bovine serum (FBS). Each cell suspension was then added to a single well of an xCELLigence E-plate 16. The proliferation rates were determined at 2-min intervals by measuring the impedance changes across the electrodes at the bottoms of the wells.

***Caspase-3 activity assay***

Human primary nasal epithelial cells (300,000) were treated with increasing concentrations of PolyP120 (up to 112 μM) or 1 mM staurosporine for 24 h. The cells were then harvested and washed with PBS and resuspended in DMEM with 10% FBS. Apoptosis was evaluated using Caspase-3 Activity Assay kits (#5723; Cell Signaling Technology), following the manufacturer instructions. Here, 4 μg total lysates were used, and the fluorescence (excitation, 380 nm; emission, 460 nm) was read using a **multimode plate readers (**PerkinElmer).

**In-vitro *infection, treatment and detection of viral RdRp gene replication in the Korean laboratory***

Vero cells (~4-6 ×10^6^) were plated into six-well plates and infected with SARS-CoV-2 viral particles (multiplicity of infection, 0.01) that were obtained from a frozen swab from a Korean patient positive for COVID-19. These experiments were performed in a B3 authorised laboratory. Uninfected Vero cells were used as the negative control of the infection. After 24 h of incubation, the infected cells were treated with the PolyPs (as sodium polyphosphate) with an indicated polymer length of 120 (PolyP120) at different concentrations: 9.38, 18.75 and 37.5 μM. Vehicle-treated cells were used as the negative treatment control. After 24 h of PolyPs treatment (i.e., 48 h from infection), the Vero cells were lysed and their RNA was extracted.

Real-time RT-PCR was performed for the specific quantitative detection of the *RdRp* gene using different target probes that were differentially labelled (AccuPower SARS-CoV-2 Real-Time RT-PCR kits; Bioneer, South Korea). These analyses were run using a PCR machine (CFX96; BioRad) under the following conditions: 25 °C for 2 min; 50 °C for 15 min; 95 °C for 2 min; 95 °C for 3 s; 55 °C for 30 s (×45 cycles).

**References**

1.	Maria Pachetti 1 2 BM, Francesca Benedetti 4, Fabiola Giudici 5, Elisabetta Mauro 3, Paola Storici 1, Claudio Masciovecchio 1, Silvia Angeletti 6, Massimo Ciccozzi 6, Robert C Gallo 7 8, Davide Zella 9 10, Rudy Ippodrino. Emerging SARS-CoV-2 mutation hot spots include a novel RNA-dependent-RNA polymerase variant. J Transl Med

2020.

2.	Kim D, Lee JY, Yang JS, Kim JW, Kim VN, Chang H. The Architecture of SARS-CoV-2 Transcriptome. Cell. 2020 May 14;181(4):914-21 e10. PubMed PMID: 32330414. Pubmed Central PMCID: 7179501.

3.	Kulaev IS. Biochemistry of inorganic polyphosphates. Reviews of physiology, biochemistry and pharmacology. 1975;73:131-58. PubMed PMID: 175427.
